# Suppression of Fatty Acid Oxidation by Thioesterase Superfamily Member 2 in Skeletal Muscle Promotes Hepatic Steatosis and Insulin Resistance

**DOI:** 10.1101/2021.04.21.440732

**Authors:** Norihiro Imai, Hayley T. Nicholls, Michele Alves-Bezerra, Yingxia Li, Anna A. Ivanova, Eric A. Ortlund, David E. Cohen

**Affiliations:** Division of Gastroenterology and Hepatology, Joan & Sanford I. Weill Department of Medicine, Weill Cornell Medical College, NY 10021 USA; Department of Biochemistry, Emory University, Atlanta, GA 30322 USA

**Keywords:** Hepatic steatosis, obesity, acyl-CoA thioesterase, fatty acid oxidation, insulin resistance

## Abstract

Thioesterase superfamily member 2 (Them2) is highly expressed in oxidative tissues where it hydrolyzes long chain fatty acyl-CoA esters to free fatty acids and CoA. Although mice globally lacking Them2 (*Them2^-/-^*) are protected against diet-induced obesity, insulin resistance and hepatic steatosis, liver-specific *Them2^-/-^* mice remain susceptible. To explore the contribution of Them2 in extrahepatic tissues, we created mice with Them2 deleted in skeletal muscle (*S-Them2^-/-^*), cardiac muscle (*C-Them2^-/-^*) or adipose tissue (*A-Them2^-/-^*). When fed a high-fat diet, *S-Them2^-/-^* but not *C-Them2^-/-^* or *A-Them2^-/-^* mice exhibited reduced weight gain. Only *S-Them2^-/-^* mice exhibited improved glucose homeostasis together with improved insulin sensitivity in skeletal muscle. Increased rates of fatty acid oxidation in skeletal muscle of *S-Them2^-/-^* mice were reflected in alterations in skeletal muscle metabolites, including short chain fatty acids, branched chain amino acids and the pentose phosphate pathway. Protection from diet-induced hepatic steatosis in *S-Them2^-/-^* mice was attributable to increased VLDL triglyceride secretion rates in support of demands of increased muscle fatty acid utilization. These results reveal a key role for skeletal muscle Them2 in the pathogenesis of diet-induced obesity, insulin resistance and hepatic steatosis.

## INTRODUCTION

Rates of obesity have increased rapidly over the past 40 years and currently affect 600 million people worldwide^1^. Obesity increases the risk of developing insulin resistance and non-alcoholic fatty liver disease (NAFLD). These common conditions contribute significantly to rising morbidity and mortality, highlighting the unmet need for effective new therapeutic targets in NAFLD.

Thioesterase superfamily member 2 (Them2; also known as acyl-CoA thioesterase (Acot) 13) catalyzes the hydrolysis of long chain fatty acyl-CoA esters into free fatty acids and CoA^2^. Them2 is abundant in the liver and primarily localized to mitochondria, where it regulates trafficking of fatty acids to oxidative versus lipid biosynthetic pathways, depending upon metabolic conditions^2^. Them2 is also expressed at high levels in other oxidative tissues, including skeletal and cardiac muscle, as well as in adipose tissue. However, its function in these tissues is not understood^3^. Mice globally lacking Them2 (*Them2^-/-^*) are protected against diet-induced obesity and hepatic steatosis, and exhibit markedly improved glucose homeostasis^4, 5^. The global absence of Them2 is also associated with reduced hepatic ER stress in the setting of overnutrition^6^. Conversely, Them2 liver-specific knockout (*L-Them2^-/-^*) mice exhibit similar body weights as controls, and are not protected against hepatic steatosis or insulin resistance in response to high-fat diet (HFD) feeding^7^. Studies in *L-Them2^-/-^* mice revealed a mechanistic role for Them2 in the hepatic trafficking of acyl-CoA molecules towards triglyceride synthesis and incorporation into VLDL particles^7^.

To explore the hypothesis that Them2 activity in extrahepatic tissues may be required for the development of insulin resistance and hepatic steatosis, we generated tissue-specific knockout mice lacking Them2 in skeletal muscle, cardiac muscle and adipose tissue. Only skeletal muscle-specific Them2 knockout mice recapitulated the protection from insulin resistance and hepatic steatosis that was observed in *Them2^-/-^* mice. Our results demonstrate that Them2 limits rates of fatty acid oxidation in skeletal muscle, and its upregulation in the setting of overnutrition promotes skeletal muscle insulin resistance and hepatic steatosis by complementary mechanisms. In addition to reducing the demand for hepatic VLDL secretion, the reduction in fatty acid oxidation in skeletal muscle decreases the production of short chain fatty acids and pentose phosphate pathway intermediates that maintain insulin sensitivity. In addition, Them2 suppresses the expression of myokines capable of modulating skeletal muscle physiology, as well as crosstalk with liver and adipose tissue. When taken together, these findings suggest that Them2 in skeletal could be targeted as therapeutic strategy in the management of NAFLD.

## RESULTS

### Them2 suppresses insulin sensitivity in skeletal muscle and limits exercise capacity in HFD-fed mice

Because skeletal muscle and heart extensively utilize fatty acids to produce ATP, we examined the potential contribution of Them2 to the regulation of insulin action in these tissues. We first confirmed that *Them2^-/-^* mice on a C57BL6J genetic background were protected against HFD-induced obesity (Figure 1-A,B) and glucose intolerance (Figure 1-C), as we have observed for a mixed genetic background^4^. In response to exogenous insulin administration (Figure 1-D), HFD-fed *Them2*^-/-^ mice exhibited a 62 % reduction in blood glucose concentrations compared with a non-significant 24 % decrease in *Them2^+/+^* mice, with an associated 2.6-fold increase in Akt phosphorylation in skeletal muscle, which together indicated improved insulin sensitivity in the absence of Them2 expression. There was a 1.7-fold increase in Them2 expression in skeletal muscle, but not in cardiac muscle, in HFD-fed compared to chow-fed mice (Figure 1-E).

**Figure 1.**
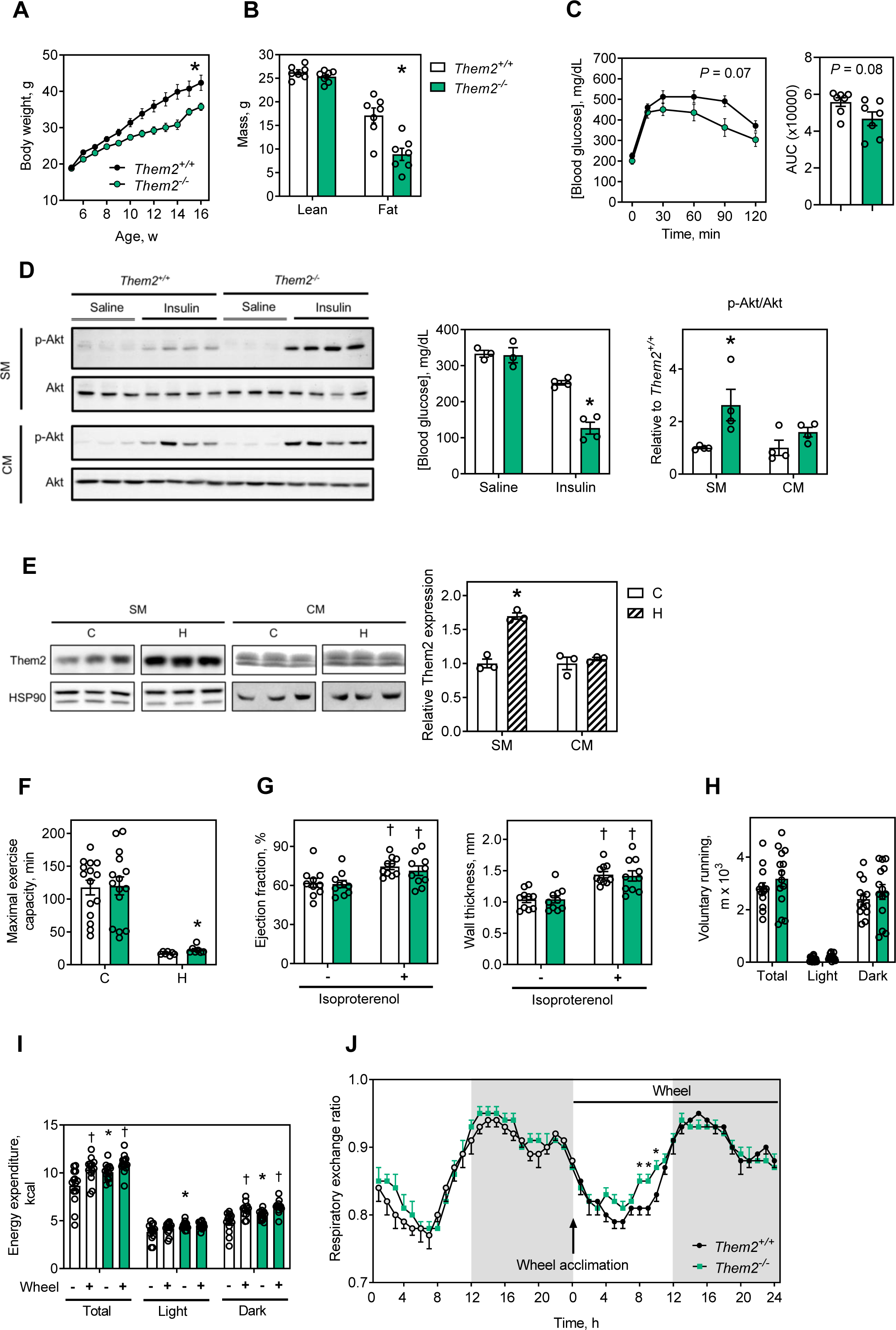
Them2 is upregulated in skeletal muscle by HFD feeding and its ablation improves skeletal muscle insulin sensitivity and exercise capacity in HFD-fed mice. Mice were fed a HFD (H) from 5-16 w. (A) Body weights and (B) Lean and fat mass (determined at 16 w of age). Tolerance tests to glucose (C) were conducted between 16 and 17 w of age. Mice were injected IP with saline or insulin (10mU/g) prior to (D) measurements of blood glucose concentrations, as well as immunoblot analysis of Akt phosphorylation in cardiac and gastrocnemius muscle tissues. (E) Expression of Them2 in gastrocnemius muscle (skeletal muscle, SM) and heart (cardiac muscle, CM). (F) Maximal exercise capacity was analyzed by graded treadmill test (n = 7-15/group). (G) Cardiac ejection fraction and left ventricle wall thickness were evaluated by echocardiography before and after isoproterenol osmotic mini pump implantation (†*P* < 0.05 before vs. after treatment). Chow-fed mice were individually housed in metabolic cages for the measurement of (H) voluntary running on an exercise wheel, (I) energy expenditure and (J) respiratory exchange ratio (n = 14/group), \**P* < 0.05 *Them2*^-/-^ vs *Them2^+^*^/+^, †*P* < 0.05 with running wheel vs without running wheel. Data are mean ± SEM. Statistical analyses were conducted using Student’s t test or two-way ANOVA with repeated-measures where appropriate.

Notwithstanding reduced body weight of *Them2*^-/-^ mice, both skeletal muscle and cardiac tissues from *Them2^-/-^* mice were proportionate in size and histologically normal (Suppl. Figure 1-A,B). Whereas skeletal muscle of *Them2^-/-^* mice exhibited no changes in mRNA expression levels of myosin heavy chains, there were modest decreases in the proportions of type I muscle fiber type relative to type II fiber type proteins (Suppl. Figure 1-C,D). *Them2^-/-^* mice also exhibited similar expression of myosin heavy chains and hypertrophy-related genes in cardiac muscle as control mice (Suppl. Figure 1-E,F). In chow-fed mice, Them2 expression did not influence in exercise capacity, as assessed by treadmill testing (Figure 1-F) and lactate production (Suppl. Figure 1-G). However, in the setting of HFD feeding, *Them2^-/-^* mice showed improved exercise capacity compared to wild type controls (Figure 1-F). We next investigated the impact of Them2 on cardiac function in response to chronic ionotropic stimulation. Whereas cardiac hypertrophy is accompanied by decreased rates of fatty acid oxidation and increased glucose utilization^8^, Them2 expression did not influence development of hypertrophy (Figure 1-G).

Metabolic monitoring revealed no differences in voluntary (wheel) running (Figure 1-H), spontaneous physical activity or food intake (data not shown). Similar to *Them2^-/-^* mice on a mixed genetic background^4^, chow-fed *Them2^-/-^* C57BL6J mice exhibited increased energy expenditure (Figure 1-I). Although no differences were found in energy expenditure during voluntary running (Figure 1-I), respiratory exchange ratios of *Them2^-/-^* mice were elevated during the light cycle after acclimation to running wheel (Figure 1-J), indicative of greater glucose utilization in skeletal muscle in the absence of Them2 expression during a period when mice were more sedentary. Taken together, these results suggest that Them2 regulates energy substrate utilization in skeletal muscle.

### Skeletal muscle expression of Them2 promotes obesity and insulin resistance in HFD-fed mice

To systematically explore contributions of extrahepatic Them2 to obesity, hepatic steatosis and insulin resistance, we created mice lacking Them2 in skeletal muscle, cardiac muscle and adipose tissues, each on a C57BL/6 genetic background (Figure 2-A). Skeletal muscle deletion of Them2 with preserved cardiac expression (*S-Them2^-/-^* mice) was accomplished using mice expressing Cre recombinase driven by the *Myf5* promoter. Because *Myf5* is expressed in brown adipose tissue (BAT) in addition to skeletal muscle, mice lacking Them2 in skeletal muscle were also deficient in BAT. The *Myh6* promoter was used to generate cardiac muscle deletion of Them2 with preserved skeletal muscle expression (*C-Them2^-/-^* mice). *A-Them2^-/-^* mice lacking Them2 in BAT and white adipose tissue (WAT), which expresses Them2 at low levels, were generated using mice expressing Cre recombinase driven by the adiponectin promoter. Only *S-Them2^-/-^* mice were protected against diet-induced overweight (Figure 2-B), excess fat accumulation (Figure 2-C), glucose intolerance (Figure 2-D), and insulin resistance (Figure 2-E), as observed for global *Them2^-/-^* mice (Figure 1-A-D). These differences were not associated with substantial alterations in expression of other skeletal muscle *Acot* or *Acsl* isoforms (Suppl. Figure 2-A,B). The possibility that this phenotype was attributable to the absence of Them2 expression in BAT was excluded by demonstrating similar weight differences (Suppl. Figure 3-A) and body compositions (Suppl. Figure 3-B) for mice housed at 30 °C, a thermoneutral temperature that suppresses BAT activity. This was further supported by the phenotypes of *A-Them2^-/-^* mice (Figure 2-B-D), which also lack Them2 expression in BAT, but were not protected against HFD-induced obesity and insulin resistance. We used a gain of function approach to confirm the metabolic contribution of skeletal muscle Them2: AAV8-Mck driven rescue of Them2 expression in muscle of global *Them2^-/-^* mice (Suppl. Figure 3-C) resulted in higher body weights than AAV8-Mck-LacZ treated *Them2^-/-^* mice (Suppl. Figure 3-D) with increases in both lean and fat mass (Suppl. Figure 3-E). Taken together, these results indicate that skeletal muscle Them2 plays a central role in the development of obesity and insulin resistance in response to HFD feeding.

**Figure 2.**
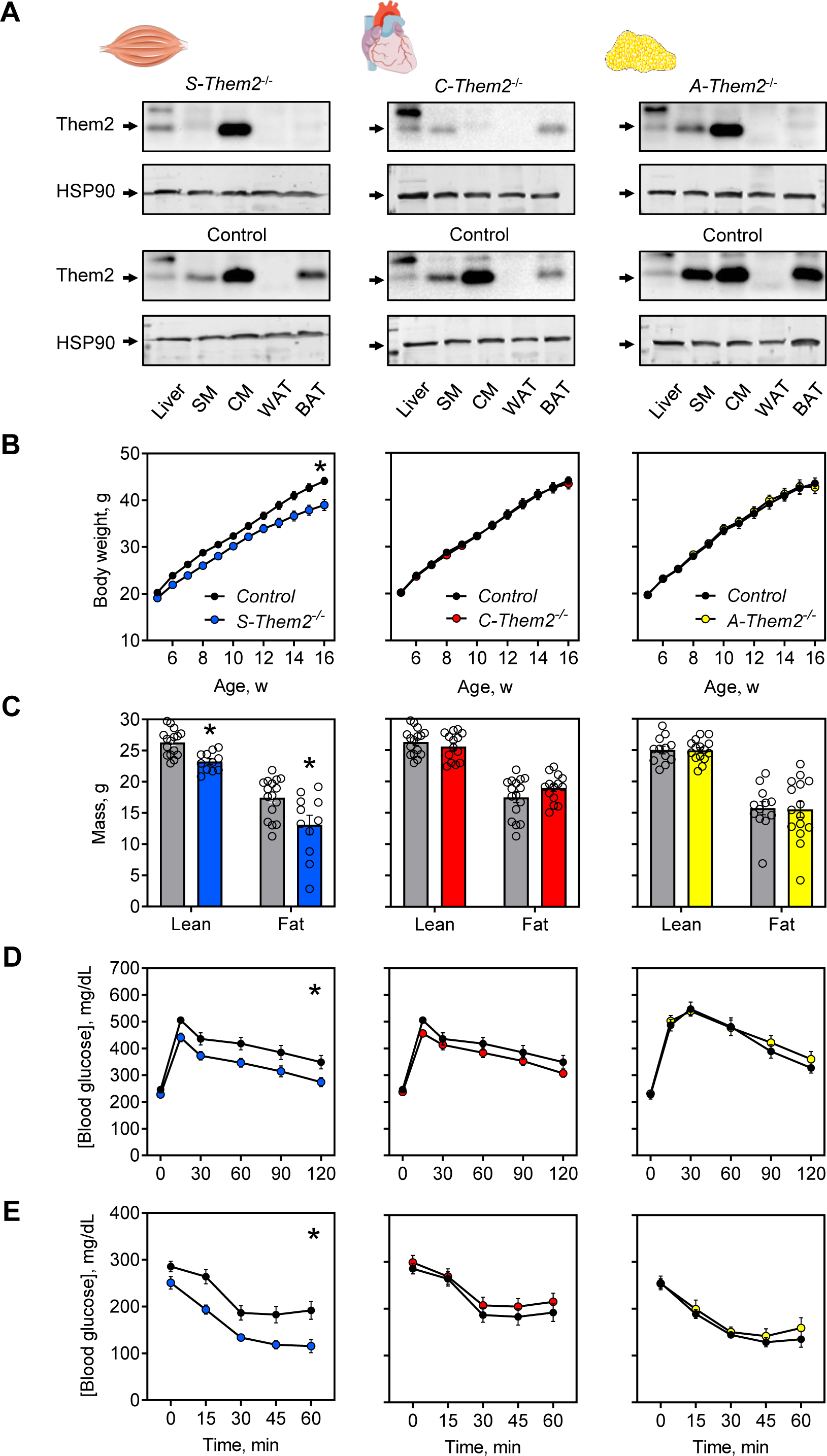
Selective deletion of Them2 in skeletal muscle reduces weight gain and improves glucose homeostasis and insulin sensitivity in HFD-fed mice. Male C57BL/6 mice with (L to R) skeletal muscle deletion of Them2 (*S-Them2*^-/-^, *Them2*^Flox/Flox^ x *Myf5-*Cre^Tg/0^); cardiac muscle deletion of Them2 (*C-Them2*^-/-^, *Them2*^Flox/Flox^ x *Myh6-Cre*^Tg/0^); and adipose tissue deletion of Them2 (*A-Them2*^-/-^, *Them2*^Flox/Flox^ x *Adipoq-Cre*^Tg/0^). (A) Immunoblot analysis of Them2 with HSP90 as a loading control. Mice were fed a HFD (H) from 5-17 w. (B) Body weights and (C) lean and fat mass (determined at 16 w). Tolerance tests to (D) glucose and (E) insulin were conducted between 16 and 17 w of age. (B-E) Data are mean ± SEM; n=6-18/group. Statistical analyses were conducted using 2-way ANOVA with repeated measures (B, D,E) or Students *t*-test (C); \**P* < 0.05 compared to control.

To investigate mechanisms of improved glucose homeostasis in HFD-fed *S-Them2^-/-^* mice, we performed hyperinsulinemic-euglycemic clamp studies. *S-Them2^-/-^* mice exhibited higher glucose infusion rates than controls (Figure 3-A), as well as increased rates of glucose uptake in skeletal muscle (Figure 3-B). By contrast, basal and clamped hepatic glucose production rates were not influenced by skeletal muscle expression of Them2 (Figure 3-C). In keeping with these findings, there were marked increases in Akt phosphorylation by 2.5-fold in skeletal muscle of HFD-fed *S-Them2^-/-^* mice compared to controls in response to acute administration of insulin (Figure 3-D). These results reveal that Them2 in skeletal muscle promotes whole body glucose intolerance by reducing skeletal muscle insulin sensitivity.

**Figure 3.**
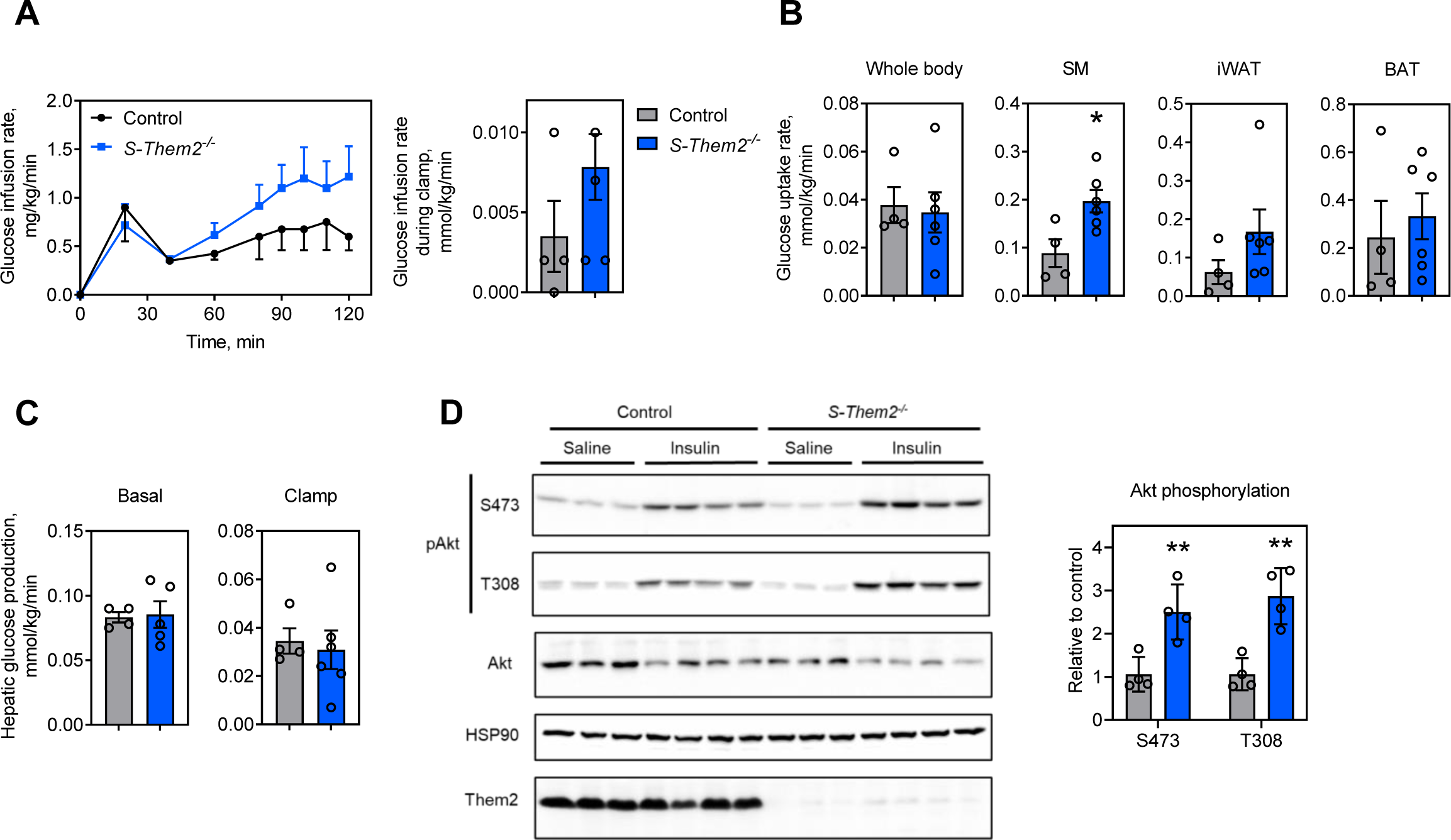
Increased glucose uptake and improved insulin sensitivity in skeletal muscle of HFD-fed *S-Them2^-/-^* mice. Hyperinsulinemic-euglycemic clamp studies were performed in mice fed a HFD for 12 w. (A) Time course of glucose infusion rates, with bar plot presenting glucose infusion rates during the clamp. (B) Glucose uptake rates were measured for whole body, gastrocnemius muscle (skeletal muscle, SM), inguinal white adipose tissue (iWAT) and brown adipose tissue (BAT). (C) Hepatic glucose production rates measured under basal and clamp conditions. (*S-Them2^-/-^*, n = 6; Control, n = 4), (D) Gastrocnemius muscle samples were collected from mice fed a HFD for 5-17 w, 10 min after i.p. injection with saline or insulin (10mU/g), and subjected to immunoblot analysis (n = 4 mice/group). **P < 0.01, *S-Them2^-/-^* vs Control. Statistical analyses were conducted using Student’s t test. Data are mean ± SEM.

### Them2 controls rates of fatty acid oxidation in skeletal muscle

We next examined the role of Them2 in the regulation of fatty acid oxidation in skeletal muscle. Rates of fatty acid oxidation in skeletal muscle homogenates were increased in both *Them2^-/-^* (Figure 4-A) and *S-Them2^-/-^* mice (Figure 4-B). Muscle strips isolated from *Them2^-/-^* mice exhibited increased oxidation rates of palmitate from endogenous (pulsed) palmitate but not from exogenous (chase) sources (Figure 4-C).

**Figure 4.**
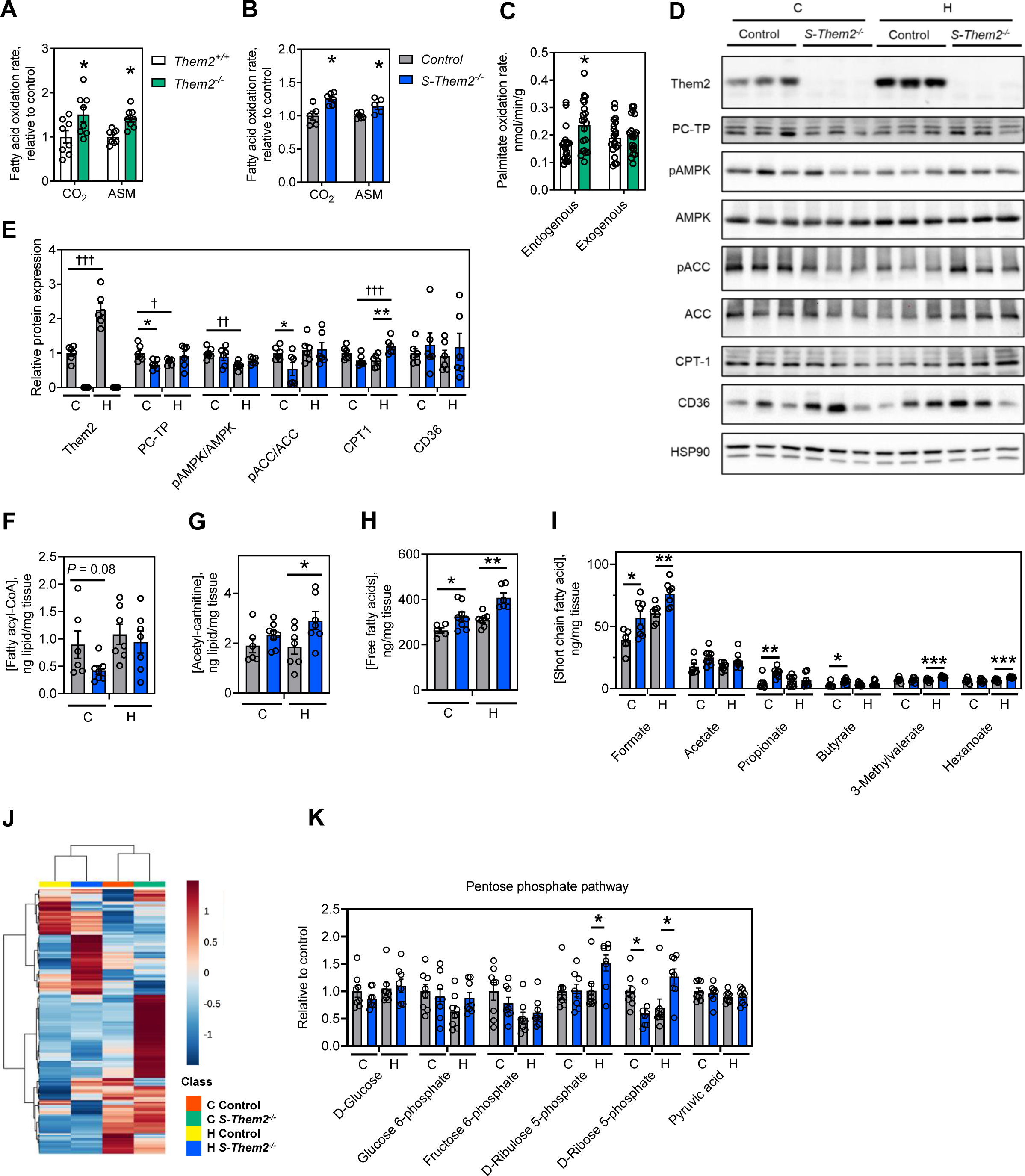
Increased oxidation of endogenous fatty acids and associated metabolic changes in skeletal muscle of *S-Them2^-/-^* mice. Fatty acid oxidation rates were assessed according to degradation rates of [^14^C]-palmitate into acid-soluble metabolites [^14^C]-labeled (ASM) and CO_2_ in homogenized gastrocnemius (skeletal) muscle from chow-fed (A) global *Them2^-/-^* and (B) *S-Them2^-/-^* mice (n = 8/group). (C) Fatty acid oxidation rates were measured in isolated soleus muscle strips from chow-fed mice using a using [^3^H]-palmitate in a pulse-chase design that differentiated between endogenous and exogenous fatty acids (n = 10/group). (D-E) Gastrocnemius samples were collected from chow-fed mice (C) or HFD (H) for 12 w. (D) Immunoblot analyses and (E) corresponding densitometry of proteins that regulate fatty acid metabolism. Skeletal muscle concentrations of (F) fatty acyl-CoAs, (G) acetyl-carnitine, (H) free fatty acids and (I) short chain fatty acids (n = 6-8/group). (J) Unsupervised clustering of data obtained by targeted profiling of 216 polar metabolites in gastrocnemius muscle. (K) Intermediates of the pentose phosphate pathway (n = 8/group). Statistical analyses were conducted using Student’s t test; \**P* < 0.05, \*\**P* < 0.01, \*\*\**P* < 0.001, *S-Them2^-/-^* vs Control. †*P* < 0.05, ††*P* < 0.01, †††*P* < 0.001, chow vs HFD. Data are mean ± SEM.

To gain mechanistic insights, we performed immunoblot analyses of gastrocnemius muscle samples to assess the expression of key proteins that control fatty acid oxidation. In contrast to chow-fed control mice, HFD-fed controls exhibited 2-fold increases in Them2 expression levels (Figure 4-D,E). Suggestive of its role in regulating Them2 expression in skeletal muscle, the Them2-interacting protein PC-TP^3^, was reduced in chow-fed *S-Them2^-/-^* mice relative to controls (Figure 4-D,E). Although phosphorylation of adenosine monophosphate-activated protein kinase (AMPK (Thr172)), which increases fatty acid oxidation rates in skeletal muscle, showed no genotype-dependent changes, chow-fed *S-Them2^-/-^* mice exhibited reduced phosphorylation of acetyl-CoA carboxylase (ACC) compared to control mice (Figure 4-D,E). Phosphorylated ACC is inactive, whereas dephosphorylated ACC enhances conversion of acetyl-CoA to malonyl-CoA, and this in turn reduces fatty acid oxidation owing to inhibition of carnitine palmitoyltransferase I (CPT1)^9^. Although CPT1 protein expression was not altered in chow-fed mice, relative mRNA expression of CPT1 was reduced in skeletal muscle of chow-fed *S-Them2^-/-^* mice (Suppl. Figure 4-A). Notwithstanding the absence of changes in ACC phosphorylation in HFD-fed mice, increased CPT1 protein expression in HFD-fed *S-Them2^-/-^* mice suggests increased mitochondrial fatty acid utilization in skeletal muscle of *S-Them2^-/-^* mice. Consistent with a lack of effect of Them2 expression on fatty acid oxidation rates of exogenous fatty acids, we observed no changes in skeletal muscle expression of CD36. To further ascertain mechanisms underlying the increased skeletal muscle fatty acid oxidation, we surveyed the expression of genes that control lipid and glucose homeostasis. The expression of these genes was largely unchanged either in chow (Suppl. Figure 4-A,B) or HFD-fed mice (Suppl. Figure 4-A,B), suggesting that increased fatty acid oxidation in *S-Them2^-/-^* mice was not compensated by fatty acid uptake or glucose utilization.

In keeping with increased rates of fatty acid oxidation, total fatty acyl-CoAs in skeletal muscle of chow-fed *S-Them2^-/-^* mice tended to decrease (Figure 4-F), but without changes in individual molecular species (Suppl. Figure 5A). The absence of a change in fatty acyl-CoAs following HFD-feeding is likely attributable to abundant cellular nutrients in the setting of overnutrition. Acetyl-carnitine forms when acetyl-CoA concentrations exceed what is required by the TCA cycle, and is essential molecules for maintaining metabolic flexibility and insulin sensitivity^10^. Skeletal muscle of *S-Them2^-/-^* mice exhibited increased acetyl-carnitine concentrations for mice fed HFD (Figure 4-G), without discernible changes in individual acyl-carnitine molecular species (Suppl. Figure 5-B). These results are consistent with increased fatty acid oxidation rates in skeletal muscle lacking Them2 even after HFD feeding. Free fatty acid concentrations were higher in skeletal muscle of *S-Them2^-/-^* mice (Figure 4-H), with increases being most pronounced in short chain molecular species fatty acids (Figure 4-I, Suppl. Figure 5-C,D), which can influence glucose homeostasis^11^.

To gain further insight into Them2 metabolic regulation in skeletal muscle, we conducted targeted metabolomics profiling for 216 predefined polar metabolites. Unsupervised clustering of differences identified both diets and genotypes (Figure 4-J). Metabolic pathway analysis identified significant alterations in *S-Them2^-/-^* mice fed either chow (Suppl. Figure 6A) or HFD (Suppl. Figure 6B). Of these, amino acid metabolism and the pentose phosphate pathway (PPP) were affected by skeletal muscle Them2 expression in mice fed both diets.

The absence of Them2 in skeletal muscle increased tissue concentrations of 9 amino acids in chow-fed mice (Suppl. Figure 7-A), whereas it decreased concentrations of 4 amino acids in HFD-fed mice (Suppl. Figure 7-B). For both diets, ablation of Them2 mainly altered branched chain amino acids. Branched-chain amino acids promote protein and glutamine synthesis in skeletal muscle, and increased levels of branched-chain amino acids is known to correlate with an improvement in glucose metabolism^12^. Therefore, increased levels of branched-chain amino acids in muscle of chow-fed *S-Them2^-/-^* mice may have contributed to protection against diet-induced insulin resistance.

Whereas most metabolites in the PPP pathway were unchanged in both chow diet and in HFD (Figure 4-K), ribose 5-phosphate was significantly reduced in chow-fed mice and both ribulose 5-phosphate and ribose 5-phosphate were increased in skeletal muscle of HFD-fed *S-Them2^-/-^* mice, suggestive of upregulation of this pathway by overnutrition in the absence of Them2. PPP generates reducing equivalents to prevent oxidative stress^13^, and its upregulation may have contributed to improved insulin sensitivity by clearing excess glucose-induced reactive oxygen species in skeletal muscle. Them2 expression in skeletal muscle did not influence activity of superoxide dismutase (Suppl. Figure 8-A) or thiobarbituric acid reactive substances (Suppl. Figure 8-B), suggesting that the ablation of Them2 did not promote oxidative stress in skeletal muscle. Taken together, these studies revealed that Them2 in skeletal muscle modulates tissue concentrations of short chain fatty acids, branched chain amino acids and PPP metabolites, each of which may play a role in the regulation of glucose homeostasis.

### Them2 limits fatty acid utilization in cultured myotubes

To assess whether Them2 alters mitochondrial fatty acid utilization in skeletal muscle, we utilized C2C12 myotubes that were transduced to overexpress mouse Them2. AAV8-Them2 transduction increased Them2 expression by 1.5-fold (Figure 5-A). Although increased Them2 expression did not alter myotube diameter or fusion index^14^, AAV8-Them2 treated myotubes exhibited greater myotube areas, suggesting Them2 expression enhances myotube elongation (Figure 5-B,C). Myotubes that overexpressed Them2 also exhibited higher rates of oxygen consumption (OCR) during both basal and maximal respiration (Figure 5-D). Of note, when palmitate was made available, there were no differences in basal or maximal respiration (Figure 5-D), with differences between bovine serum albumin (BSA) treatment alone and palmitate plus BSA exposure signifying oxidation of extracellular palmitate. As evidenced by higher OCR values following treatment with BSA alone, AAV8-Them2 transduced myotubes utilized less extracellular palmitic acid (Figure 5-E).

**Figure 5.**
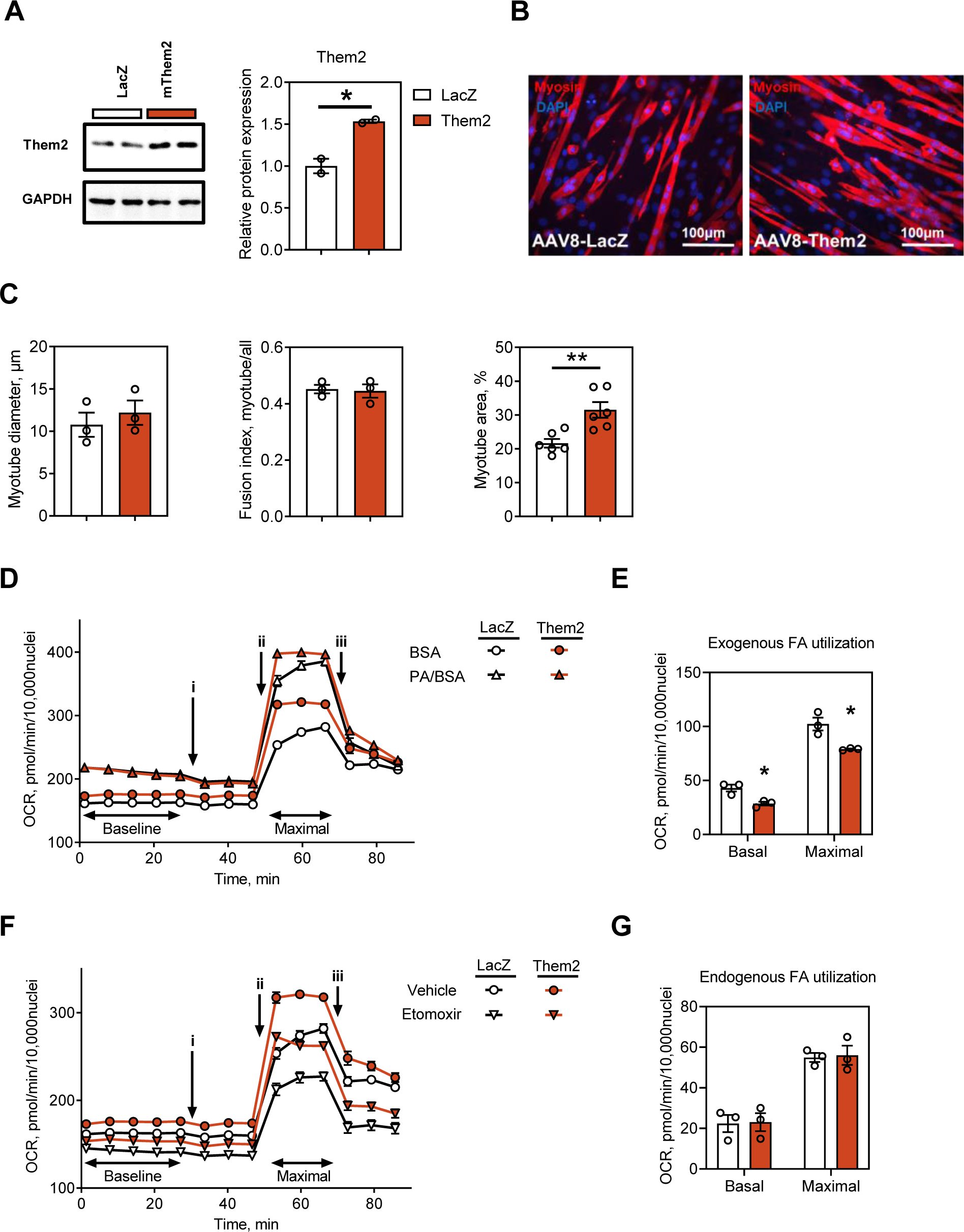
Them2 regulated myocellular fatty acid utilization. Cultured C2C12 myotubes were transduced with 1 × 10^5^ GC AAV8-Mck-LacZ or AAV8-Mck-Them2. (A) Immunoblot analysis of Them2, with GAPDH used to control for unequal loading. (B) Representative images of transduced C2C12 myotubes. Cells were fixed and immunostained with anti-myosin antibody (red) to reveal sarcomeres. Cell nuclei were stained with DAPI (blue). (C) Myotube area, diameter and fusion index were evaluated using Image J software. Oxygen consumption rates (OCR) of AAV8 transduced C2C12 myotubes were measured by Seahorse XF96e following to the sequential injections of (i) oligomycin, (ii) FCCP and (iii) rotenone/antimycin A. (E) OCR values were in the presence of bovine serum albumin (BSA) treated and palmitic acid (PA) plus BSA. (F) Utilization of exogenous fatty acids (FA) was determined by subtracting OCR values in the presence of BSA alone from PA/BSA. (F) OCR values were measured in the presence of the CPT-1 inhibitor etomoxir or vehicle (DMSO). (G) Utilization of endogenous FA was determined by subtracting OCR in the presence of etomoxir from vehicle. Data represent combined results from 3 independent experiments. Bar plots present FA utilization during the baseline (basal) and the 3 time points following the addition of FCCP (maximal). Statistical analyses were conducted using Student’s t test; \**P* < 0.05, \*\**P* < 0.01. Data are mean ± SEM.

A reduction in OCR after etomoxir treatment under substrate limited conditions provides a measure of intracellular fatty acid oxidation (Figure 5-F), which was not influenced by Them2 expression (Figure 5-G). Taken together, these results suggest that Them2 regulates extracellular fatty acid utilization through its regulation of the intracellular fatty acid pool that is available for oxidation.

### Skeletal muscle Them2 expression enables HFD-fed mice to exceed a body weight threshold value for hepatic steatosis and metabolic dysfunction

In human beings, hepatic steatosis is largely a function of adipose tissue insulin resistance and serves an important barometer of metabolic dysfunction, which becomes fully established at a threshold level of 5.5% liver fat^15^. Although HFD-fed mice are widely utilized in studies of NAFLD, a threshold value for pathogenic intrahepatic triglyceride accumulation and its relationship to body composition has not been systematically explored. To enable accurate weight-based comparisons between *S-Them2^-/-^* and control mice, 87 C57BL6J male mice were fed a HFD for 12 w. Body weights exhibited a normal distribution ranging from 31.1 g to 52.2 g (mean ± SEM; 42.3 ± 0.5 g; median 42.5g) (Figure 6-A). Both lean and fat mass increased linearly as functions of body weight (Figure 6-B), and fat mass was a linear function of lean mass (Figure 6-C). By contrast, liver weight exhibited two very distinct linear relationships to body weight (Figure 6-D): Up to a body weight threshold of 41.6 g (corresponding to fat mass of 15.6 g), there was a relatively flat slope of liver weight versus body weight (0.031 g liver/g BW). In excess of 41.6 g, the slope was 4.3-fold steeper (0.134 g liver/g BW). Additionally, there was a similar biphasic dependence of liver weight on fat (Figure 6-E) but not lean weight of mice (Figure 6-F).

**Figure 6.**
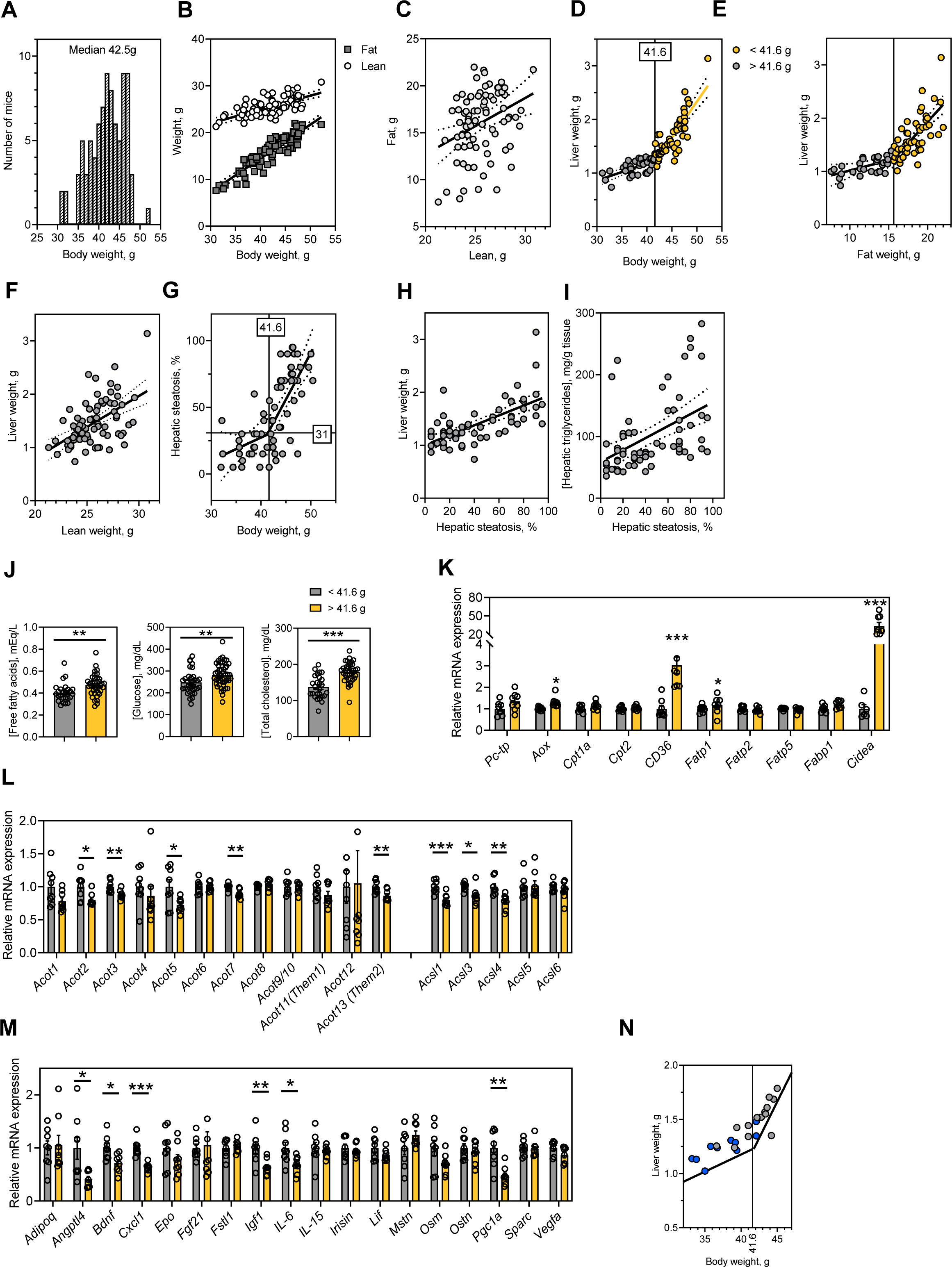
Skeletal muscle Them2 expression enables HFD-fed mice to exceed a threshold value for accelerated hepatic steatosis and metabolic dysfunction. C57BL/6J mice were fed a HFD for 12 w. (A) Body weight distribution (n = 84) and (B) fat and lean weights. (C) Linear relationship between fat and lean weights. (D-E) Biphasic relationships between liver weight and (D) body weight and (E) fat weight (n=64). Threshold value of 41.6 g body weight was determined by segmental linear regression. (F) Linear relationship between liver weight and lean weight. (G) Biphasic relationship between hepatic steatosis determines histologically and body weight. (H) Linear relationship of liver weight to hepatic steatosis. (I) Linear relationship of hepatic triglycerides concentrations to hepatic steatosis. (n=64). (J) Plasma concentrations of free fatty acids, glucose and total cholesterol. (K) Hepatic mRNA expression levels of selected genes that govern the metabolism of fatty acids and lipid droplets (n = 8/group). Relative mRNA expression in skeletal muscle (L) of genes that regulate fatty acid utilization and (M) myokines. (N) Body and liver weights of HFD-fed *S-Them2^-/-^* and control mice (n = 10-14/group) relative to the biphasic linear regression derived for HFD-fed C57BL/6J mice (panel D). Statistical analyses were conducted using Student’s t test; \**P* < 0.05, \*\**P* < 0.01, \*\*\**P* < 0.001, body weight < 41.6 g vs body weight > 41.6 g. Data are mean ± SEM. Abbreviations: *Pctp*: phosphatidylcholine transfer protein, *Aox*: alternative oxidase, *CPT-1*: carnitine palmitoyltransferase I, *Fatp*: fatty acid transport protein, *Fabp*: fatty acid binding protein, *Cidea*: cell death activator, *Acot*: acyl-CoA thioesterase, *Acsl*: acyl-CoA synthetase long chain family member, *Adipoq*: adiponectin, *Angptl4*: angiopoietin-like 4, *Bdnf*: brain derived neurotrophic factor, *CXCL1*: chemokine ligand 1, *Epo*: erythropoietin, *Fgf21*: fibroblast growth factor 21, *Fstl1*: follistatin-like 1, *Igf1*: insulin like growth factor 1, *IL-6*: interleukin-6, *IL-15*: interleukin-15, *Lif*: leukemia inhibitory factor, *Mstn*: myostatin, *Osm*: oncostatin M, *Ostn*: osteocrin, *Pgc1a*: *Pparg* coactivator 1 alpha, *Sparc*: secreted protein acidic and rich in cysteine, *Vegfa*: vascular endothelial growth factor A.

Hepatic steatosis exhibited a biphasic relationship to body weight, with threshold body weight of 41.6 g corresponded to 31% hepatic steatosis (Figure 6-G). By contrast, both liver weights (Figure 6-H) and hepatic triglyceride concentrations (Figure 6-I) were linearly correlated with histologic steatosis. The body weight threshold of 41.6 g also discriminated plasma concentrations of free fatty acids, glucose and cholesterol (Figure 6-J), as well as hepatic mRNA expression levels of specific genes that govern the metabolism of fatty acids (*Aox* and *CD36*) and lipid droplets (*Cidea*) (Figure 6-K).

Indicative that the 41.6 g weight threshold for hepatic steatosis reflected changes in skeletal muscle dysfunction, expression of genes that mediate fatty acid activation (*Acsls*) and deactivation (*Acots*) were significantly altered in skeletal muscle of mice exceeding this body weight (Figure 6-L). Additionally, five myokine genes were downregulated above this weight threshold, as was *Pgc1a*, which controls mitochondrial biogenesis in skeletal muscle (Figure 6-M). Taken together, these findings reveal two distinct phases in the development of hepatic steatosis and metabolic dysfunction in HFD-fed C57BL/6J mice with a threshold of 31% at a body weight of 41.6 g. After 12 w of HFD-feeding, body weights of most (80%) of *S-Them2^-/-^* mice fell below the 41.6 g threshold, compared with 36 % of controls (Figure 6-N).

In keeping with reduced weight and improved glucose homeostasis, HFD-fed *S-Them2^-/-^* but not *L-Them2^-/-^*^7^ or *C-Them2^-/-^* mice were also protected against hepatic steatosis, as evidence by histopathology (Figure 7-A) and hepatic triglyceride concentrations (Figure 7-B), as was previously observed in *Them2^-/-^* mice^4^. Pyruvate tolerance tests revealed reduced hepatic glucose production only in *S-Them2^-/-^* mice (Figure 7-C), as was also observed in global *Them2^-/-^* mice^4^. There were no differences in plasma activities of aspartate aminotransferase, alanine aminotransferase or creatine kinase or in insulin concentrations (Suppl. Figure 9-A,B). Of note, for mice under the 41.6 g body weight threshold, hepatic triglyceride concentrations were also reduced in HFD-fed *S-Them2^-/-^* mice compared to controls (Figure 7-D). These findings suggested that metabolic improvements in *S-Them2^-/-^* mice were in part attributable to reduced body weight gain because most mice lacking Them2 did not achieve a body weight threshold for the development of more severe hepatic steatosis, but importantly, were also due in part to expression of skeletal muscle Them2 *per se*.

**Figure 7.**
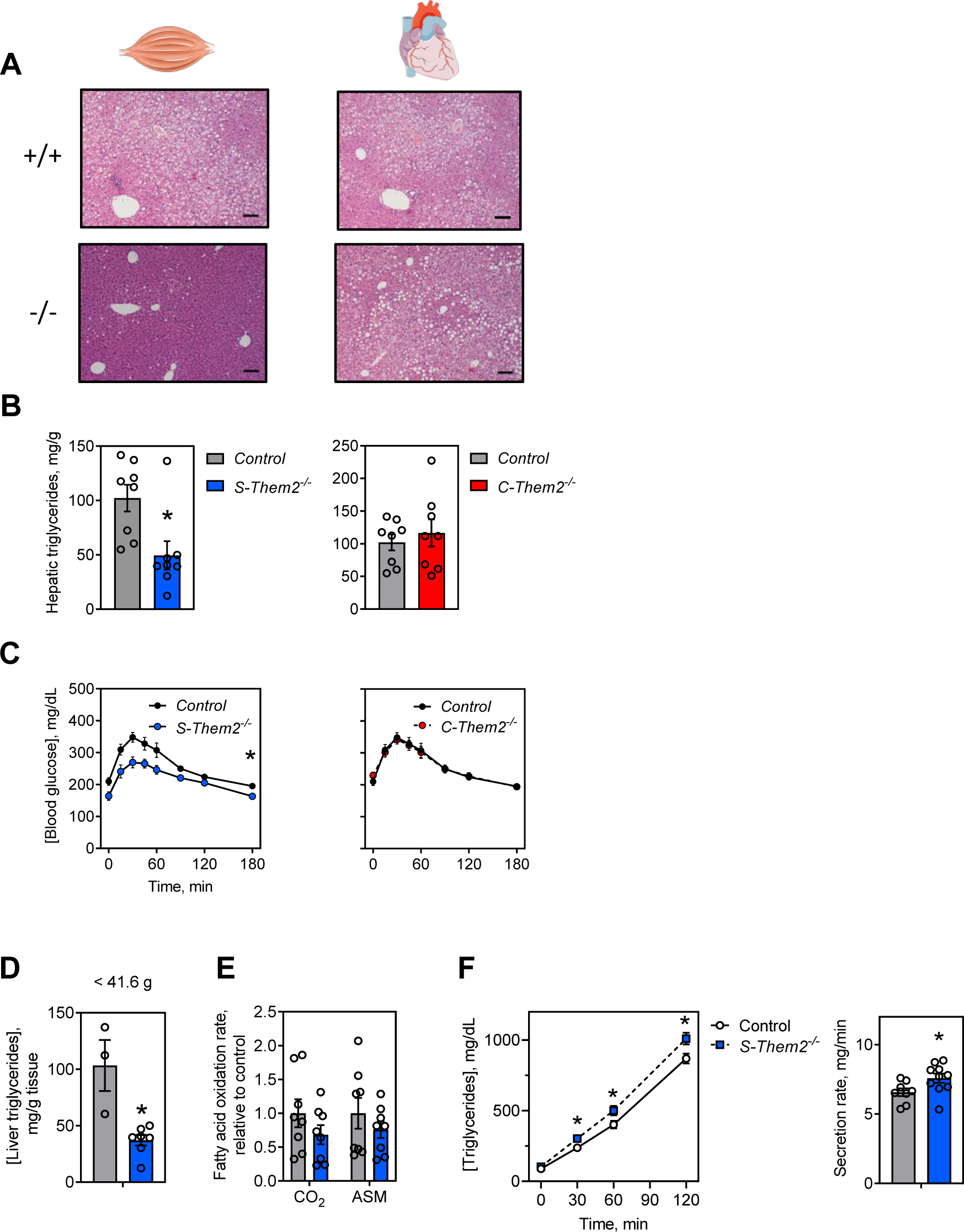
Them2 promotes hepatic steatosis by suppressing VLDL secretion. (L to R) Mice with skeletal muscle (*S-Them2^-/-^*) and cardiac muscle (*C-Them2^-/-^*) Them2 deletion were fed a HFD for 12 w and then livers were analyzed for (A) hepatic steatosis by H&E staining (bars represent 100µm) and (B) hepatic triglyceride concentrations. (C) Pyruvate tolerance tests conducted in HFD-fed mice at 16 w of age. Data are mean ± SEM (n=6-12/group). Statistical analyses were conducted by Students t-test (B) or 2-way ANOVA with repeated measures (C); \**P* < 0.05 vs control. Data are presented as mean ± SEM (D) Hepatic triglyceride concentrations of HFD-fed *S-Them2^-/-^* and control mice with body weights of less than 41.6 g (n = 3-7/group). (E) Fatty acid oxidation rates in livers of HFD-fed *S-Them2^-/-^* mice (n = 8/group). (F) Rates of triglyceride hepatic secretion in HFD-fed mice (n = 8-10/group). Statistical analyses were conducted using Student’s t test or two-way ANOVA with repeated-measures where appropriate. \**P* < 0.05, *S-Them2^-/-^* vs Control. Data are mean ± SEM.

### Skeletal muscle Them2 expression promotes hepatic steatosis by suppressing VLDL secretion

We next explored the mechanisms underlying skeletal muscle Them2-mediated development of hepatic steatosis. We first examined evidence for increased rates of hepatic fatty acid oxidation. Fasting concentrations of ß-hydroxybutyrate were reduced in HFD-fed control mice compared with chow-fed controls, but this was not observed for *S-Them2^-/-^* mice, in which levels remained unchanged by HFD-feeding and were significantly higher than in HFD-fed control mice (Suppl. Figure 9-C). This suggested that the absence of Them2 in skeletal muscle might be associated with increase rates of fatty acid oxidation in liver. However, direct measurement of hepatic fatty acid oxidation in HFD-fed *S-Them2^-/-^* mice did not reveal increases (Figure 7-E), suggesting that differences in ß-hydroxybutyrate may instead have reflected Them2-mediated changes in rates of ketone body utilization in skeletal muscle. Chow but not HFD-fed *S-Them2^-/-^* mice exhibited increased fasting plasma concentrations of free fatty acids (Suppl. Figure 9-D), which argues against reduced hepatic uptake of fatty acids as a mechanism by which *S-Them2^-/-^* mice were protected against steatosis. Plasma concentrations of triglycerides, total and free cholesterol, as well as phospholipids were unaffected by skeletal muscle Them2 expression in either chow or HFD-fed mice (Suppl. Figure 9-D), and despite dramatic reductions in hepatic steatosis, RNAseq analysis of liver identified only a limited number of genes with altered expression (Suppl. Figure 10-A). Among these was the hepatic steatosis marker *Cidea*, which was markedly downregulated, suggesting reductions in triglyceride storage. This was also implicated by pathway analysis, which identified increased degradation of hepatic triglycerides as the most upregulated pathway (Suppl. Figure 10-B). In keeping with this possibility, *S-Them2^-/-^* mice exhibited increased rates of VLDL triglyceride secretion (Figure 7-F). Taken together, these findings suggest that increased demand by skeletal muscle for fatty acids in the form of secreted hepatic triglycerides may have explained the reduced steady state concentrations in liver.

### Muscle Them2 mediates crosstalk between skeletal muscle and the liver

Lastly, we examined evidence of regulatory cross-talk between muscle and liver that may have also contributed to protection against hepatic steatosis and metabolic dysfunction in *S-Them2^-/-^* mice. Conditioned medium from C2C12 myotubes transduced to overexpress Them2, suppressed basal and maximal OCR values in Hepa1-6 cells (Figure 8-A). We next assessed for Them2-dependent expression of myokines in skeletal muscle. Ablation of Them2 altered myokine gene expression depending on dietary conditions (Figure 8-B). Overnight fasting induced substantial changes in myokine genes expression in *S-Them2^-/-^* mice (Figure 8-B). Skeletal muscle of chow-fed *S-Them2^-/-^* mice induced expression of insulin like growth factor 1 (*Igf-1*) and interleukin 15 (*IL-15*), both of which regulate muscle mass^16, 17^, whereas *IL-15* also decreases plasma triglycerides and adipose tissue mass^18^. In HFD-fed mice, *S-Them2^-/-^* mice exhibited modest increases in chemokine ligand 1 (*CXCL1*), which regulates angiogenesis and fatty acid metabolism^19^. Independent of diet, there was consistent down-regulation in *osteoclin* which is upregulated by palmitic acid and promotes insulin resistance in muscle^20–22^. Because skeletal muscle of HFD-fed *S-Them2^-/-^* mice is protected against excess body weight gain, these alterations in myokine genes might have been secondary to changes in body weight. However, differences in relative mRNA expression were still observed in weight matched HFD-fed mice, as defined by body weights that fell below the threshold of 41.6 g (Figure 8-C): *myostatin* and *osteoclin* were downregulated in skeletal muscle of *S-Them2^-/-^* mice compared to control mice. These results suggested that Them2-mediated control of skeletal muscle myokine gene expression may contribute to crosstalk among skeletal muscle, adipose tissue and the liver (Figure 8-D).

**Figure 8.**
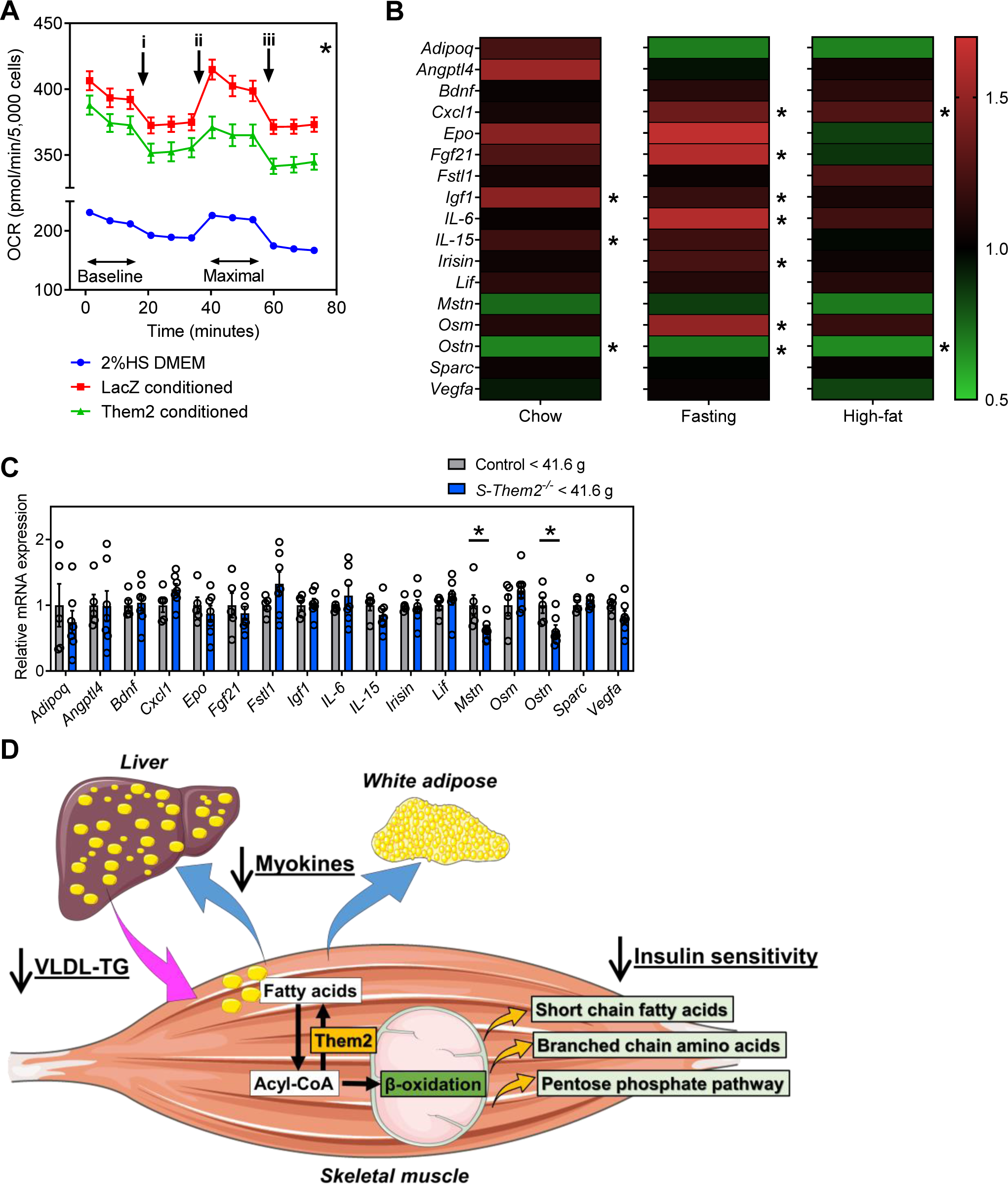
Skeletal muscle Them2 mediates crosstalk between muscle and the liver. (A) Rates of oxygen consumption (OCR) of Hepa1-6 cells cultured with conditioned medium. Sequential additions of oligomycin (i), FCCP (ii) and rotenone/antimycin A (iii) are indicated. Conditioned medium was collected from C2C12 myotubes reconstituted with AAV-LacZ or AAV-Them2. Data represent combined results from 3 independent experiments. (B) Relative mRNA expression of myokine genes in soleus (skeletal) muscle dissected from mice fed chow diet or a HFD for 12 w. Myokine genes were also analyzed in chow-fed mice after 18 h fasting. Heat map represents relative values to controls for the same diet. (C) Relative mRNA expression of myokine genes in body weight matched (<41.6g) mice fed HFD for 12 w (n = 5-7/ group). \**P* < 0.05, *S-Them2^-/-^* vs body weight matched (<41.6 g) control. (D) Schematic summary of metabolic role of Them2 in skeletal muscle and its influence on the development of hepatic steatosis. Them2 in skeletal muscle decreases dependence on fatty acid oxidation to produce ATP, therefore utilizing less circulating free fatty acids. Them2 limits fatty acid oxidation in skeletal muscle consequently altering the production of short chain fatty acids, branched chain amino acids and pentose phosphate pathway. These alterations are most evident in the setting of over nutrition and limit insulin sensitivity in skeletal muscle. Them2 also regulates muscle-to-organ crosstalk by multiple myokine genes depending on dietary condition, therefore Them2 may coordinate metabolism of fatty acids in skeletal muscle, adipose tissue and the liver. Statistical analyses were conducted using Student’s t test (B,C) or two-way ANOVA with repeated-measures (A). Data are mean ± SEM.

## DISCUSSION

This study identified a key role for skeletal muscle Them2 in the regulation of skeletal muscle metabolism, as well as in the pathogenesis of HFD-induced insulin resistance and hepatic steatosis in mice. Them2 limits rates of fatty acid oxidation by skeletal muscle, which in turn reduces rates of insulin-mediated glucose disposal by reducing insulin sensitizing metabolites and myokines. This also reduced demand for exogenous fatty acids, leading to the development of hepatic steatosis.

We previously demonstrated that *Them2^-/-^* mice exhibited reduced plasma free fatty acid concentrations, as well as reduced hepatic steatosis and glucose production^5, 23^. However, these effects were not recapitulated in *L-Them2^-/-^* mice, suggesting that extrahepatic Them2 expression is required in order to recapitulate the phenotype in the setting of overnutrition^7^. In both mice and humans, skeletal and cardiac muscles substantially contribute to basal calorie consumption. Skeletal plays critical roles in energy metabolism, glucose uptake and locomotion^24^. The metabolic activity of the heart is a major determinant of the body’s oxygen consumption, predominantly utilizing long chain fatty acids as substrates for energy production^8^. Consequently, glucose and lipid metabolism in skeletal muscle and cardiac muscle contribute substantially to whole body energy consumption.

Whereas Them2 is most highly expressed in heart, its deletion from cardiac muscle did result in discernable differences in organ function or whole body metabolism. To preserve function, cardiac muscle has extraordinary metabolic flexibility that enable it to adapt to changes in substrate availability^8^. For this reason, we speculate that alterations in this specific enzyme of long chain fatty acid metabolism would not cause obvious functional or metabolic consequences in *C-Them2^-/-^* mice. This notion is supported by our observation that Them2 expression in heart of control mice was unaffected by HFD feeding. By contrast, HFD feeding robustly upregulated Them2 expression in skeletal muscle, suggesting a pathogenic role in skeletal muscle mechanical dysfunction and alteration in metabolism.

As evidenced by reduced rates of glucose uptake during clamp studies and diminished insulin-stimulated Akt phosphorylation, Them2 contributed to the development of skeletal muscle insulin resistance, presumably as a reflection of reduced rates of fatty acid oxidation^25^. Previous studies have demonstrated that the accumulation of excess free fatty acids and their derivatives including long chain acyl-CoAs in skeletal muscle are key determinants of insulin resistance^26^. Although the absence of a fatty acyl-CoA thioesterase might have been expected to increase skeletal muscle concentrations fatty acyl-CoAs and decrease concentrations of free fatty acids, we observed the opposite in *S-Them2^-/-^* mice. However, the preferred substrate of Them2 is long chain fatty acyl-CoA, and the observed increase in free fatty acid concentrations was mainly attributable to increases in short chain fatty acids. Short chain fatty acids are predominantly generated by colonic bacteria and are metabolized by enterocytes and hepatocytes. Short chain fatty acids are also formed in peroxisomes during shortening of very long chain fatty acyl-CoAs that can be hydrolyzed by peroxisomal *Acots* and released into the cytosol^27^. We speculate, increased rates of mitochondrial ß-oxidation in skeletal muscle of *S-Them2^-/-^* mice were also accompanied by increases in peroxisomal oxidation, which resulted in the production of more short chain fatty acids. The reduction in skeletal muscle fatty acyl-CoA concentrations most likely reflects their increased rates of oxidation, as evidence by the reciprocal rise in acetyl-carnitine concentrations.

Short chain fatty acids influence muscle fatty acid oxidation, glucose uptake and insulin sensitivity via both direct and indirect mechanisms^11, 28^. Of note, acetate and butyrate improve skeletal muscle glucose metabolism and insulin sensitivity^11^. When taken together with observed increases in branched chain amino acids and PPP metabolites^12, 13^, these data suggest that elevated concentrations of short chain fatty acids in skeletal muscle of *S-Them2^-/-^* mice contributed to the improved insulin sensitivity observed the setting of overnutrition.

Although the reduction in lean mass of *S-Them2^-/-^* mice at the thermoneutrality was similar as at room temperature, there was less reduction in fat mass. This may have been because Them2 in BAT also plays a role in HFD-induced obesity via adaptive thermogenesis^5^. Because fat mass is tightly correlated with lean mass in HFD-fed mice, we further postulate that reduction in lean body mass due to loss of skeletal muscle Them2 also exerted metabolic control through associated decreases in adipose tissue. In human NAFLD, hepatic steatosis is strongly influenced by relatively modest changes in total body weight: An 11% loss in body weight was associated with a 52 % reduction in hepatic triglycerides, owing largely to the restoration of insulin sensitivity^29^. Interestingly, these observations in humans are of a similar magnitude to our findings in HFD-fed mice, whereby at 16 w of age, *S-Them2^-/-^* mice exhibited a 12 % reduction in BW (Figure 2-B), which was associated with a 52 % reduction in hepatic triglyceride concentrations. However, unlike observations in humans^15^, C57BL6J mice exhibited remarkable accelerations in hepatic triglyceride accumulation at a discrete threshold of body weight (41.6 g). Moreover, fat weight is tightly correlated with lean weight in HFD-fed mice. Hence, we concluded that reduced hepatic steatosis in *S-Them2^-/-^* mice is attributable in part to the reduced body weight gain such that mice rarely exceeded body weights of 41.6 g. Nevertheless, hepatic triglyceride concentrations were still reduced in HFD-fed *S-Them2^-/-^* mice compared to HFD-fed control mice weighing less than 41.6 g revealing that Them2 expression *per se* is sufficient to regulate hepatic lipid metabolism.

A potential mechanism whereby skeletal muscle Them2 directly controls hepatic steatosis is organ crosstalk through the secretion of myokines, which are bioactive proteins that regulate organ function by autocrine, paracrine and endocrine actions^30^. Deletion of Them2 in skeletal muscle induced expression of several myokines. Skeletal muscle of chow-fed *S-Them2^-/-^* mice showed significantly higher gene expressions of *Igf-1* and *IL-15*, which regulate muscle hypertrophy^16, 17^. *IL-15* also decreases lipid deposition in pre-adipocytes and reduces white adipose tissue mass^18^. In the setting of overnutrition, there was also a modest increase in skeletal muscle expression of *CXCL1* in *S-Them2^-/-^* mice. *CXCL1* is the functional homologue to human *IL-8*^31^, and it plays a role in angiogenesis and neutrophil chemoattractant activity^32^. Increased expression of *CXCL1* attenuates diet induced obesity in mice through enhanced fatty acid oxidation in muscle^19^. *Osteoclin* (synonym: *musclin*), which was downregulated in skeletal muscle of *S-Them2^-/-^* mice, is a myokine that is controlled by nutritional status and promotes insulin resistance. *Musclin* expression inhibits glucose uptake and glycogen synthesis^20^, and *Musclin* mRNA is regulated by the forkhead box O1 transcription factor (*FOXO1*) downstream of *PI3K/Akt* pathway in response to insulin action^22^.

Our findings demonstrate a central role for skeletal muscle Them2 in the control of nutrient metabolism and in the pathogenesis of diet-induced insulin resistance and hepatic steatosis. The inhibition of Them2 activity in muscle could offer an attractive strategy for the management of obesity and nonalcoholic fatty liver disease.

## MATERIALS AND METHODS

### Animals and diets

Mixed background global Them2 knockout mice were backcrossed 20 generations with C56BL/6J^4, 5^. Wild type C56BL/6J mice were obtained from the Jackson Laboratory, Bar Harbor, ME, USA (Stock #000664). Tissue specific knockout mice were created by a LoxP/Cre system^7^. C56BL6 background mice with Them2 flanked by two LoxP sites (*Them2*^flox/flox^) were crossed with mice expressing Cre recombinase driven by the Myf5 promoter to generate skeletal muscle specific deletion of *Them2^-/-^* (*S-Them2^-/-^*), the Myh6 promoter to generate cardiac muscle specific deletion of *Them2^-/-^* (*C-Them2^-/-^*) and the Adipoq promoter was used to generate adipose tissue specific deletion of *Them2^-/-^* (*A-Them2^-/-^*) respectively (The Jackson Laboratory, Myf5-cre; Stock #007893, Myh6-cre; Stock #011038, Adipoq-cre; Stock #010803)^33–35^. These mice were viable and displayed no apparent developmental abnormalities. The presence of Cre allele was determined by PCR analysis using the primers specified by the Jackson Laboratory. Mice were housed in a barrier facility on a 12 h light/dark cycle with free access to water and diet. Male mice were weaned at 4 w of age and fed chow (PicoLab Rodent Diet 20; LabDiet, St. Louis, MO, USA). Alternatively, 5 w old male mice were fed a HFD (D12492: protein 20 % kcal, fat 60 % kcal, carbohydrate 20 % kcal, energy density 5.21 kcal/g; Research Diets Inc.,

New Brunswick, NJ, USA) for 12 w. Following 6 h fasting (9:00 AM to 3:00 PM) or 18 h fasting (6:00 PM to 12:00 PM), 17 w old mice were euthanized and plasma was collected by cardiac puncture. Tissues were harvested for immediate use or snap frozen in liquid nitrogen and stored at -80 °C. Mice were monitored by daily health status observation by technicians supported by veterinary care. Housing and husbandry were conducted in facilities with a sentinel colony health monitoring program and strict biosecurity measures to prevent, detect, and eradicate adventitious infections. Animal use and euthanasia protocols were approved by the Institutional Animal Care and Use Committee of Weill Cornell Medical College.

### Adeno-associated virus-mediated Them2 expression in skeletal muscle

Adeno-associated virus (AAV) vectors used in this study were generated by Vigene Biosciences, Rockville, Maryland, USA. Vectors are based on the AAV8 genome and transgenes are driven by the muscle creatine kinase (Mck) promoter. The mouse Them2 gene (gene ID: 55856) was packaged with Flag as AAV8-Mck-Flag-Them2 and a LacZ gene was used to generate AAV8-Mck-LacZ as a control. Mice were anesthetized with 2% isoflurane vapors prior to injection to ensure no movement. Both hindlimbs were shaved and sterilized with isopropanol swab sticks. AAVs (1-5x10^11^ particles in 30μl saline) were injected to the gastrocnemius muscle by intramuscular injection using 30-gauge insulin syringe.^36^ Mice were then return to home cages for 12 w to allow for recombinant protein expression.

### Maximal exercise capacity

Mice were acclimated to treadmill running tracks incrementally for 5 d starting from 10 min at 0.3 km/h to 60 min at 1.2 km/h. To determine maximal exercise capacity, treadmill speed was started at 0.3 km/h and increased 0.3 km/h every 3 min until the speed reached 1.2 km/h. The speed was kept constant (1.2 km/h) until mice reached exhaustion. Blood lactate levels were measured from the tail tip before and after 30 min treadmill running at 1.2 km/h using lactate pro2 LT-1730 (ARKRAY, Kyoto, Japan).

### Cardiac function

Cardiac function was evaluated by echocardiography as response to 2 w chronic isoproterenol infusion as previously described^37–39^. In brief, 10 w old mice were implanted with ALZET osmotic minipumps Model 1002 (DURECT Corporation, Cupertino, CA, USA) and infused with isoproterenol at 30 mg/kg/day for 2 w. Echocardiography was carried out at 10 w (pre) and 12 w (post) to monitor cardiac hemodynamics using a Vevo 3100 (FUJIFILM Visual sonics, Toronto, Canada). M mode was used to measure left ventricle ejection fraction and wall thickness. Each mouse was anesthetized with inhaled isoflurane (2%).

### Metabolic monitoring

Mice were housed in individual cages for 2 w for acclimation prior to metabolic monitoring^40^. Mice were then housed in temperature-controlled cabinets with a 12 h light/dark cycle and monitored using the Promethion Metabolic Screening System (Sable Systems International, North Las Vegas, NV). Rates of O_2_ consumption (VO_2_) and CO_2_ production (VCO_2_) were determined at 5 min intervals. Values of respiratory exchange ratio (RER) were calculated as VCO_2_/VO_2_. After 2 d acclimation, metabolic parameters were recorded over 24 h with or without a running wheel inside the cage. Physical activities and voluntary running on wheel were determined according to beam breaks within a grid of photosensors built outside the cages. Energy expenditure was calculated by indirect calorimetry^41^ and adjusted by ANCOVA using VassarStats (www.vassarstats.net) to adjust for differences in lean body mass using lean body composition determined by magnetic resonance imaging EchoMRI (EchoMRI, Houston, TX).

### Tolerance Tests

Glucose tolerance tests, insulin tolerance tests, and pyruvate tolerance tests were performed with minor modifications^7, 42^. In brief, mice were fasted 6 h for glucose tolerance tests, 4 h for insulin tolerance tests, and overnight (16 h) for pyruvate tolerance tests. A drop of blood from the tail tip was subjected to glucose measurement at baseline and at regular intervals using GE 100 Blood Glucose Monitor (General Electric, Ontario, CA). Glucose solution was administered by intraperitoneal injection with 1 g/kg for global *Them2^-/-^* mice and their controls and with 1.5 g/kg for *A-Them2^-/-^* mice and their littermate controls, or orally using gavage needle with fixed dose at 40 mg for *C-Them2^-/-^* mice, *S-Them2^-/-^* mice and their littermate controls. Insulin solution was administered by intraperitoneal injection with 0.75 U/kg for *A-Them2^-/-^* mice, *C-Them2^-/-^* mice, *S-Them2^-/-^* mice and littermate controls. Pyruvate solution was administered by orally using a gavage needle with 2 g/kg for *C-Them2^-/-^* mice, *S-Them2^-/-^* mice and littermate controls.

### Hyperinsulinemic-euglycemic clamp studies

Hyperinsulinemic-euglycemic clamp studies were performed at Animal Physiology Core (Albert Einstein College of Medicine, New York, NY, USA) as described privously^43^. Briefly, vascular catheters were inserted, 8 d prior to the clamps. Hyperglycemic-euglycemic clamp studies were performed in unrestrained mice. Mice were fasted for 5 h prior to the study. The clamp protocol consisted of a 90 min tracer (5-uCi bolus of [3-^3^H]-glucose followed by a 0.05 uCi/min infusion) equilibration period followed by a 120 min experimental period. A blood sample (10 ul) was taken for the assessment of basal glucose and insulin levels and glucose turnover. The insulin clamp was begun with a primed-continuous infusion of human insulin (300 mU/kg bolus followed by 5 mU/kg/min 1; Novolin; Novo Nordisk, Princeton, NJ). The [3-^3^H]-glucose infusion was increased to 0.1 uCi/min for the remainder of the experiment. Euglycemia (150 mg/dl) was maintained during clamps by measuring blood glucose every 10 min and infusing 45% dextrose as necessary. Blood samples (60–200 µl) were taken and processed to determine glucose specific activity. All blood samples were resuspended in heparinized saline and infused back to mice to maintain hematocrit. Plasma samples were collected to determine glucose levels and specific activities of [3-^3^H]-glucose. Tissue samples were stored at −80°C for glucose uptake analyses^43^.

### Insulin signaling

Mice were fasted for 6 h during the light cycle (9:00 AM to 3:00 PM). Tissues were collected 10 min following intraperitoneal insulin (10 U/kg) injection or saline injection as a control. Insulin signaling was evaluated by immunoblot analysis for phosphorylation of Akt^44^.

### Quantitative PCR

Relative mRNA expression was determined by quantitative PCR (qPCR) using SYBR Green Real-Time PCR Master Mix (Applied Biosystems, Foster City, CA, USA). Total RNA was isolated from soleus muscle (10 µg) and apex of ventricle (20 µg) using RNeasy fibrous tissue mini kit (QIAGEN, Hilden, Germany) following manufacturer’s instructions. cDNA was synthesized with a High-Capacity cDNA Reverse transcription Kit (Applied Biosystems, Foster City, CA, USA). Equal amount of mRNA samples were subjected to qPCR using QuantStudio 6 Flex Real-Time PCR System (Applied Biosystems, Foster City, CA, USA) in a 384 plate. Genes were analyzed relative to cyclophilin A. The nucleotide sequences of oligonucleotides used for qPCR are presented in Suppl. Table 2.

### Immunoblot analysis

Immunoblot analyses were performed by standard techniques^7^. Briefly, tissues or cells were homogenized in a RIPA buffer containing cOmplete Protease Inhibitor Cocktail and PhosphoSTOP Phosphate Inhibitor Cocktail Tablets (Roche, Indianapolis, IN, USA) using a Bead Ruptor 24 Elite bead mill homogenizer (Omni International, Kennesaw, GA, USA). Protein concentrations were determined by using a BCA reagent (Thermo Fisher Scientific, Springfield Township, NJ). Equal amounts of protein samples were separated by SDS-PAGE and transferred to nitrocellulose membranes for immunoblot analysis. Membranes were incubated with primary antibodies (Suppl. Table 1) overnight and probed with respective secondary antibodies (Dako-Agilent, Santa Clara, CA, USA) for 1 h. Bands were then developed using enhanced chemiluminescence (SuperSignal West DURA, Thermo Fisher Scientific, Springfield Township, NJ, USA) and imaged by ChemiDoc XRS+ (BioRad, Hercules, CA, USA).

### Electrophoresis for myosin heavy chains

Frozen soleus muscle strips were homogenized in a lysis buffer containing 0.5M NaCl, 20mM NaPPi, 50mM Tris, 1mM EDTA and 1mM DTT, adjusted pH 6.8 using a Bead Ruptor 24 Elite bead mill homogenizer (Omni International, Kennesaw, GA, USA). Supernatants from 10 min centrifugation at 2500 x g were diluted 1:1 with glycerol. Samples are then suspended in 1:1 in a Laemmli sample buffer and incubate at room temperature for 10 min and then heated (70°C) for 10 min. Myosin heavy chains were separated by SDS-PAGE in a 6% acrylamide gel containing 30% glycerol over 24 h. Bands were visualized by Coomassie Brilliant Blue staining^45^.

### Histopathology

Freshly harvested tissues from 17 w-old mice were immersed in 10% neutralized formaldehyde for 2 d. Following to a paraffin embedding, sectioned tissues were stained with Hematoxylin and Eosin by the Laboratory of Comparative Pathology, Memorial Sloan Kettering Cancer Center, NY, USA. Slides were visualized using an Eclipse Ti microscope (Nikon, Tokyo, Japan).

### Tissue triglyceride concentrations

Lipids were extracted from frozen specimens with a mixture of chloroform/methanol (2:1) using Folch’s method as previously described^7^. Concentrations of triglycerides were assayed enzymatically (FUJIFILM Wako Diagnostics, Mountain View, CA).

### Plasma assays

Enzymatic assay kits were used to measure plasma concentrations of triglycerides, free fatty acids, total cholesterol, free cholesterol, phospholipid (FUJIFILM Wako Diagnostics, Mountain View, CA), ß-hydroxybutyrate (Stanbio Laboratories, Boerne, TX), aspartate aminotransferase, alanine aminotransferase (Infinity AST/ALT, Thermo Scientific, Waltham, MA, USA) and creatine kinase (Stanbio CK Liqui-UV Test, EKF Diagnostics, Cardiff, UK). Plasma concentrations of insulin were measured by using an ELISA kit (Crystal Chem, Downers Grove, IL), according to the manufacturer’s protocol.

### Rates of fatty acid oxidation

Rates of fatty acid oxidation in muscle tissues were measured by degradation of ^14^C-palmitate (American Radiolabeled Chemicals; St. Louis, MO, USA) into ^14^C acid soluble metabolites (ASM) and ^14^C-labeled CO_2_^7, 46^. Briefly, gastrocnemius muscle strips were collected from 6 h fasted mice. Tissues were kept on ice no longer than 30 min. Muscle tissues were minced and homogenized in a Dounce homogenizer followed by centrifugation for 10 min at 420 x g. Supernatants were transferred to microtubes containing 0.4 μCi ^14^C-palmitate / 500 μM palmitate conjugated with 0.7 % fatty acid-free BSA and incubated at 37 °C for 30 min. ^14^C-labeled CO_2_ produced by TCA cycle was captured onto filter paper soaked with 1 M NaOH and ^14^C-labeled ASM were separated with 1 M perchloric acid. ^14^C-labeled CO_2_ in the filter paper and ^14^C-labeled ASM in the supernatant were dissolved in Ecoscint H (National Diagnostics; Atlanta, GA, USA) and were counted using a liquid scintillation counter (Beckman Coulter; Danvers, MA).

Fatty acid oxidation in skeletal muscle was also evaluated by applying pulse chase design^47^. Soleus muscle strips were freshly dissected and incubated in preoxygenated HBSS with 10 mM HEPES, Ca and Mg supplemented 2 mM pyruvate, 4 % fatty acid free BSA and 1 mM palmitic acid adjusted pH 7.4. Isolated soleus muscles were then transferred into pulse buffer (HBSS with 10 mM HEPES, Ca and Mg supplemented 2 mM pyruvate, 4% fatty acid free BSA plus 2 uCi of 9,10 ^3^H-palmitate). After washing following the pulse incubation, the muscle strips with preoxygenated with incubation buffer in the absence of ^3^H-palmitate. ^3^H-palmitate pulsed muscles were then transferred to the tubes containing chase buffer (buffer: HBSS with 10 mM HEPES, Ca and Mg supplemented 2 mM pyruvate, 4 % fatty acid free BSA plus 0.5 uCi/ml of ^14^C-palmitate) and a ^14^CO_2_ trap. ^3^H_2_O produced from the endogenous oxidation of [9,10-^3^H] palmitate was separated from chase buffer with a mixture of chloroform/methanol (2:1). Gaseous ^14^CO_2_ produced from the exogenous oxidation of [1-^14^C] palmitate was measured by transferring the chase buffer to 1 M H_2_SO_4_ and a paper disc with 1 M NaOH. Liberated ^14^CO_2_ was trapped in the paper disc. Samples were dissolved in Ecoscint H (National Diagnostics; Atlanta, GA, USA) and counted using a liquid scintillation counter (Beckman Coulter; Danvers, MA).

### Hepatic VLDL triglyceride secretion rates

Following a 12 h fast, the lipoprotein lipase inhibitor Tyloxapol (500 mg/kg of body weight) (Sigma-Aldrich) was administered through retro-orbital injection.^7^ Tail tip blood samples (25 μL) were collected into microtubes before Tyloxapol injection and at regular intervals for up to 2 hours. Serum triglyceride concentrations were determined using the enzymatic assay described above. Rates of hepatic triglyceride secretion were calculated from the time-dependent linear increases in serum triglyceride concentrations.

### Lipidomics

Fatty acids, fatty acyl-CoAs and acyl-carnitines were analyzed by LC MS/MS via Multiple Reaction Monitoring on an ExionLC AC system coupled to triple quadrupole mass spectrometer QTRAP5500 (ABSciex, Framingham, MA, USA) at the Emory Integrated Lipidomics Core (Emory University, Atlanta, GA, USA)^7^. Briefly, quadriceps muscle samples were homogenized using Bead Ruptor Elite (Omni International, Kennesaw, GA, USA). Extracted samples were derivatized with 3-Nitrophenylhydrazine for fatty acids and n-butanol containing 5 % acetyl chloride for carnitines. The resuspended samples were subjected to LC MS/MS analysis. Data processing, peak determination, and peak area integration were performed using the MultiQuant 3.0.2 software (AB Sciex, USA).

### Polar metabolite profiling

Predefined 216 targeted polar metabolites were analyzed at Proteomics & Metabolomics Core Facility (Weill Cornell Medicine, New York, NY, USA). Briefly, gastrocnemius muscle tissues were homogenized in 80 % methanol and lyophilized. Measurements were normalized by tissue weight as described priviously^48^. Metabolomics pathway analysis and clustering analysis were performed using Metabo Analyst ver.4.0 (https://www.metaboanalyst.ca/home.xhtml).

### RNA sequencing

RNA sequencing was performed at Genomics Resources Core Facility (Weill Cornell Medicine, New York, NY, USA). Briefly, RNA samples from liver tissues were extracted by RNeasy Fibrous Tissue Mini Kit and RNeasy Mini Kit following to the manufacturer’s instruction (QIAGEN, Venlo, Netherlands). After quality check by electrophoresis, RNA sequencing was performed at 50 million reads per sample using Nova Seq 6000 (illumina, San Diego, CA, USA).

### Cell culture

C2C12 myoblasts (ATCC #CRL-1772) were maintained in high glucose DMEM with 10 % bovine serum and 1 % penicillin-streptomycin. To induce myotubes differentiation, C2C12 myoblasts were plated at 25,000 cells/cm^2^. Once reaching 100 % confluency, media was changed to DMEM based differentiation medium with 2 % HS and 1 % penicillin-streptomycin. Differentiation medium was changed 24 h later and every 48 h thereafter. Myotubes were transduced with AAV8-Mck-Flag-Them2 or AAV8-Mck-LacZ at 1×10^5^ genome copies after 4 d in differentiation medium. Experiments were conducted on d 6 of differentiation.

### Mitochondrial respiration

Cellular metabolic flux was determined by measuring oxygen consumption rate (OCR) in Hepa1-6 cells (ATCC #CRL-1830) with conditioned medium obtained from C2C12 myotubes reconstituted with AAVs. OCR values were measured using a Seahorse XF^e^ 96 extracellular flux analyzer (Seahorse Bioscience; North Billerica, MA, USA). OCR values were measured under basal conditions and following exposure to oligomycin, carbonyl cyanide-p-trifluoromethoxyphenylhydrazone (FCCP), rotenone and antimycin A respectively. After retrieving the plate from Seahorse instrument, live cell nuclei were stained with NucRed™ Live 647 ReadyProbes™ Reagent (Thermo Fisher Scientific, Waltham, MA, USA) following to the manufacturer’s instruction. Live cell nuclei were counted in each whole well images which obtained by Spectramax i3x (Molecular Devices, San Jose, CA, USA). Normalized OCR values were calculated according to the numbers of live cell nuclei in each well.

### Statistical Analyses

Data are presented as mean values with error bars representing SEM. Statistical significance was determined by using two-tailed unpaired Student’s t-tests when two groups were compared. Correlations were evaluated by Pearson’s correlation coefficient analysis. Threshold values were determined by segmental linear regression. Multiple group comparisons were performed using two-way ANOVA. Differences were considered significant for P < 0.05 (GraphPad Prism 8, GraphPad Software, La Jolla, CA, USA).

## Acknowledgements

The authors thank Dr. Scott Rodeo, Hospital for Special Surgery for assistance with treadmill experiments. The Weill Cornell Medicine Metabolic Phenotyping Core and the Emory Integrated Lipidomics Core are subsidized by Weill Cornell Medicine and the Emory University School of Medicine respectively.

---

**Supplemental Figure 1.**
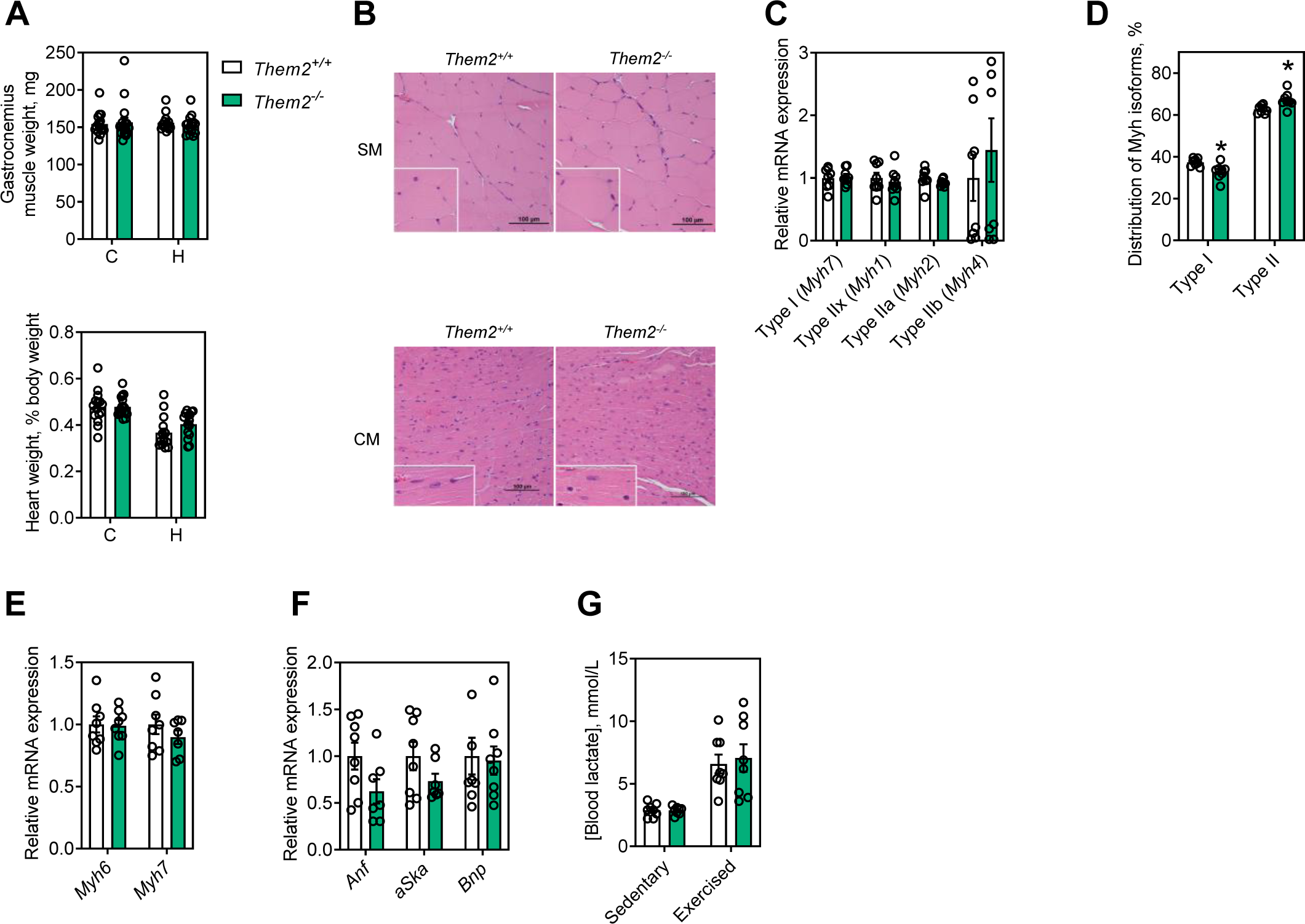
Ablation of Them2 does not promote muscle atrophy or hypertrophy. (A) Weights of cardiac and gastrocnemius (skeletal) muscle from mice fed chow (C) or HFD (H) for 12 w (n = 15-17/group, C57BL6J control mice; *Them2^+/+^*, global Them2 knockout mice; *Them2^-/-^*). (B) Histology of skeletal (gastrocnemius) muscle (SM) and cardiac muscle (CM). (C) Relative mRNA expression of myosin heavy chain genes in soleus muscle. (D) Relative mRNA expression of myosin heavy chain and cardiac hypertrophy genes in ventricle (n = 8/group). (E) Myosin heavy chain isoforms in soleus muscle determined by electrophoresis separation (n = 8/group, \**P* < 0.05, *Them2^-/-^* vs *Them2^+/+^*). (F) Blood lactate concentrations were measured before (sedentary) and 30 min after treadmill running (exercised mice) (n =8/group). Statistical analyses were conducted using Student’s t test. Data are mean ± SEM. Abbreviations: *Myh*, myosin heavy chain; *Anf*, atrial natriuretic factor; *aSka*, α-skeletal actin; *Bnp*, brain natriuretic peptide.

**Supplemental Figure 2.**
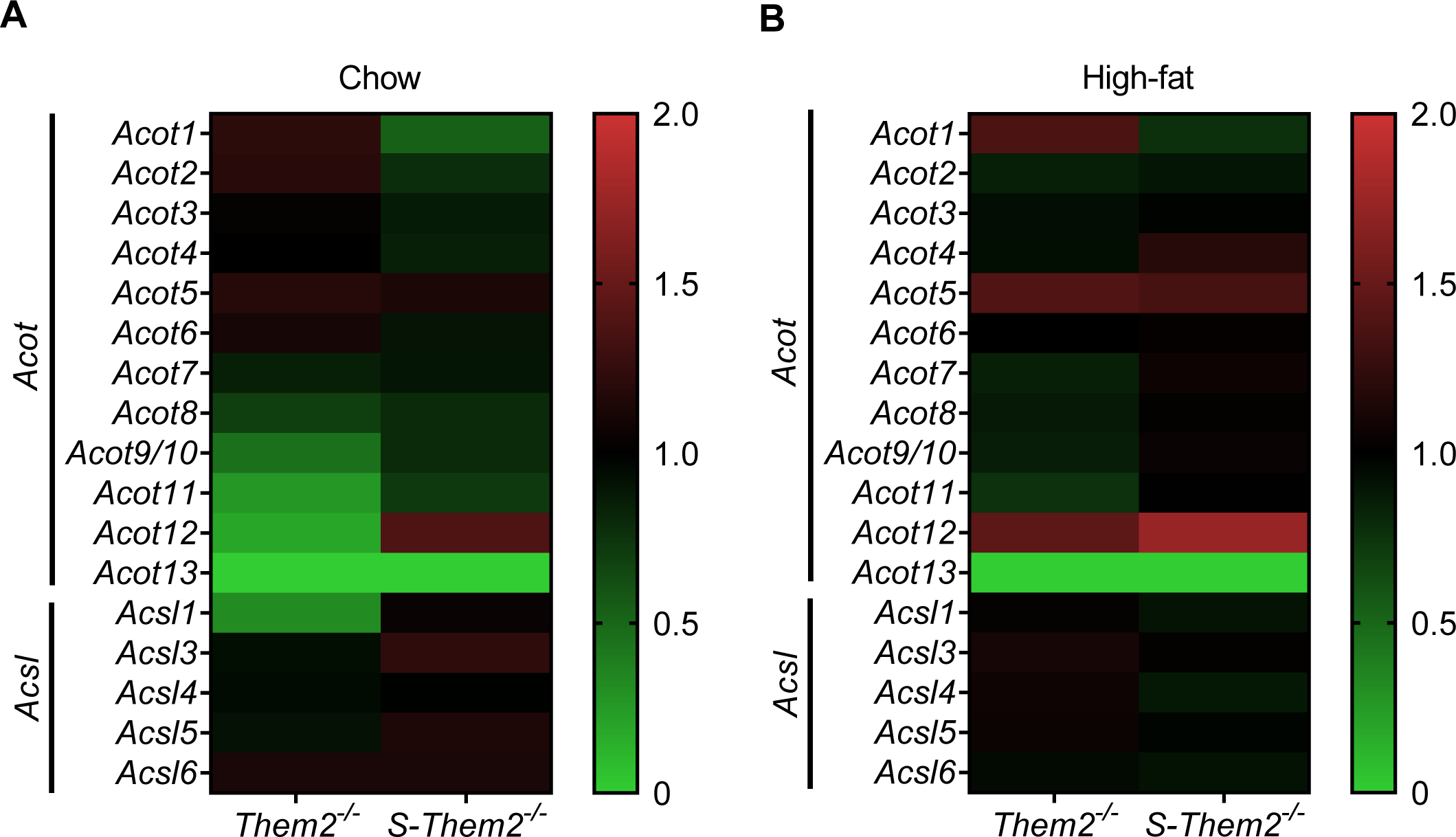
Effect of skeletal muscle Them2 on the gene expression of *Acot* and *Acsl* genes. (A) Relative mRNA expression levels of *Acot* and *Acsl* genes in skeletal muscle of chow-fed mice (n = 8/group). (B) Relative mRNA expression of *Acot* and *Acsl* genes in skeletal muscle of HFD-fed mice (n = 8/group). Heat maps represent relative gene expression compared to C57BL6 for *Them2^-/-^* and floxed control for *S-Them2^-/-^*.

**Supplemental Figure 3.**
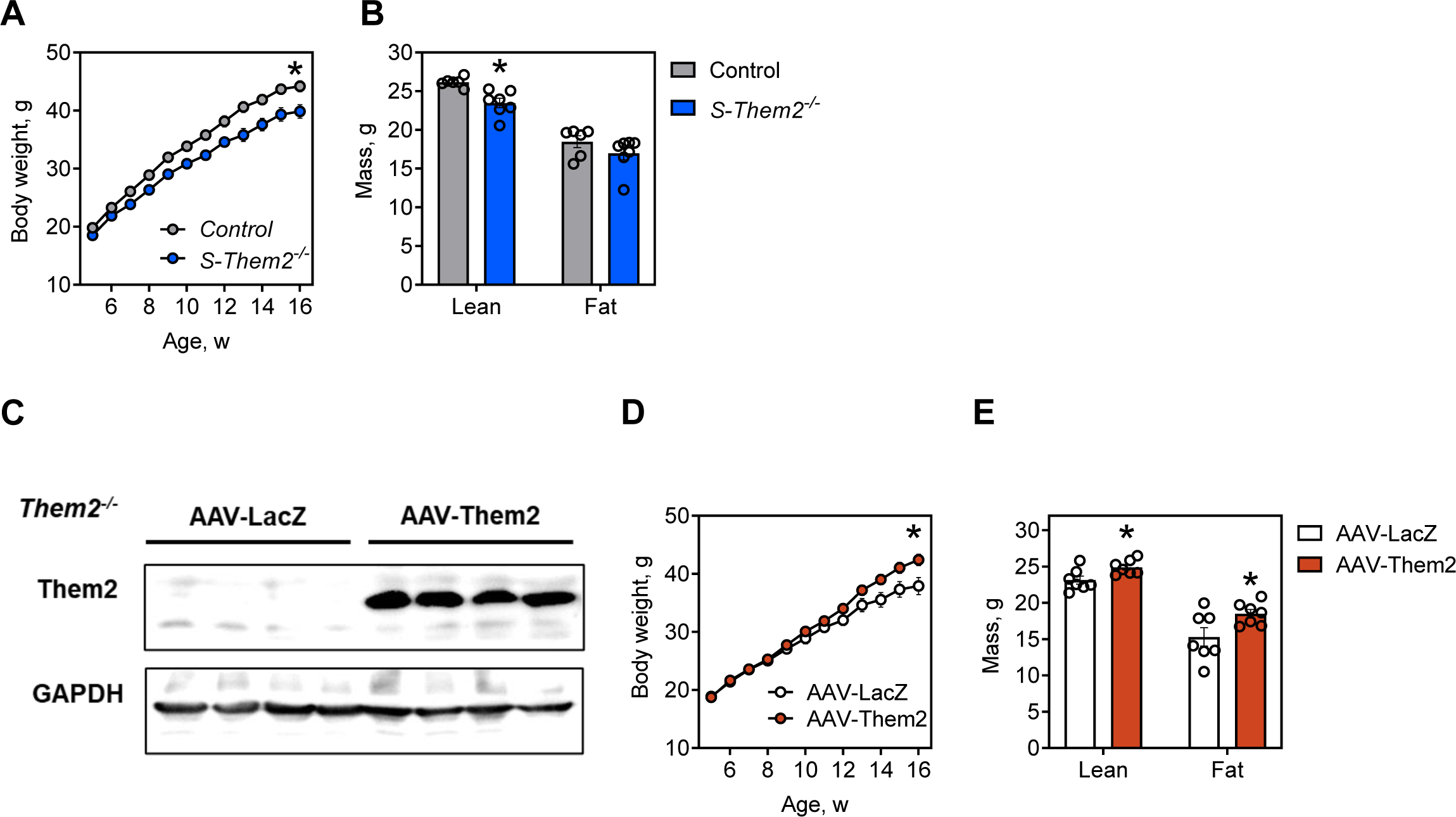
Skeletal muscle expression of Them2 mediates excess weight gain in mice fed HFD. (A) Body weights of HFD-fed mice housed at 30 °C (n=6-7/group). \**P* < 0.05 vs control. (B) Body composition of mice fed a HFD for 12 w while housed at 30 °C (n=6-7/group). Statistical comparisons were conducted using Student’s t-test; \**P* < 0.05 vs control. (C) Immunoblot analysis of Them2 in gastrocnemius muscle with GAPDH as a control for unequal loading. (D) Body weights of HFD-fed *Them2^-/-^* mice following treatment with AAV carrying Them2 or LacZ as a control (n=7/group). P < 0.05; AAV-LacZ vs AAV-Them2. (E) Body composition of HFD-fed mice. Mice were maintained at 30 ℃ (n=7/group). *P < 0.05; AAV-LacZ vs AAV-Them2. Statistical analyses were conducted using Student’s t test or two-way ANOVA with repeated-measures where appropriate. Data are mean ± SEM.

**Supplemental Figure 4.**
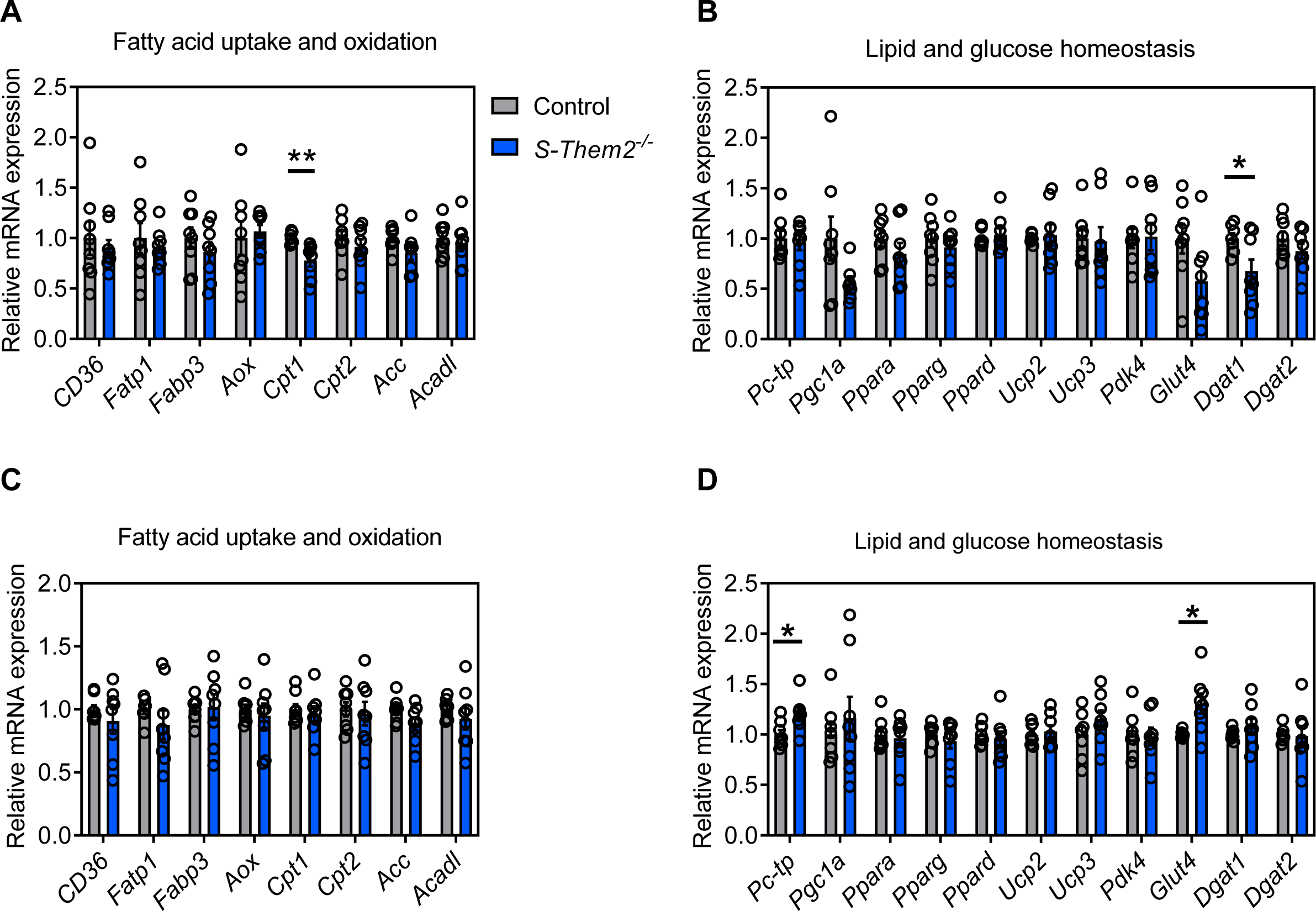
Effect of skeletal muscle Them2 expression on the expression of genes that control lipid and glucose homeostasis. Mice were (A,B) fed chow or (C,D) HFD for 12 w. Relative mRNA expression of genes in skeletal muscle that (A,C) regulate fatty acid uptake and oxidation and (B,D) lipid and glucose metabolism (n = 8/group). Statistical analyses were conducted using Student’s t test; \**P* < 0.05, \*\**P* < 0.01 vs control. Data are mean ± SEM.

**Supplemental Figure 5.**
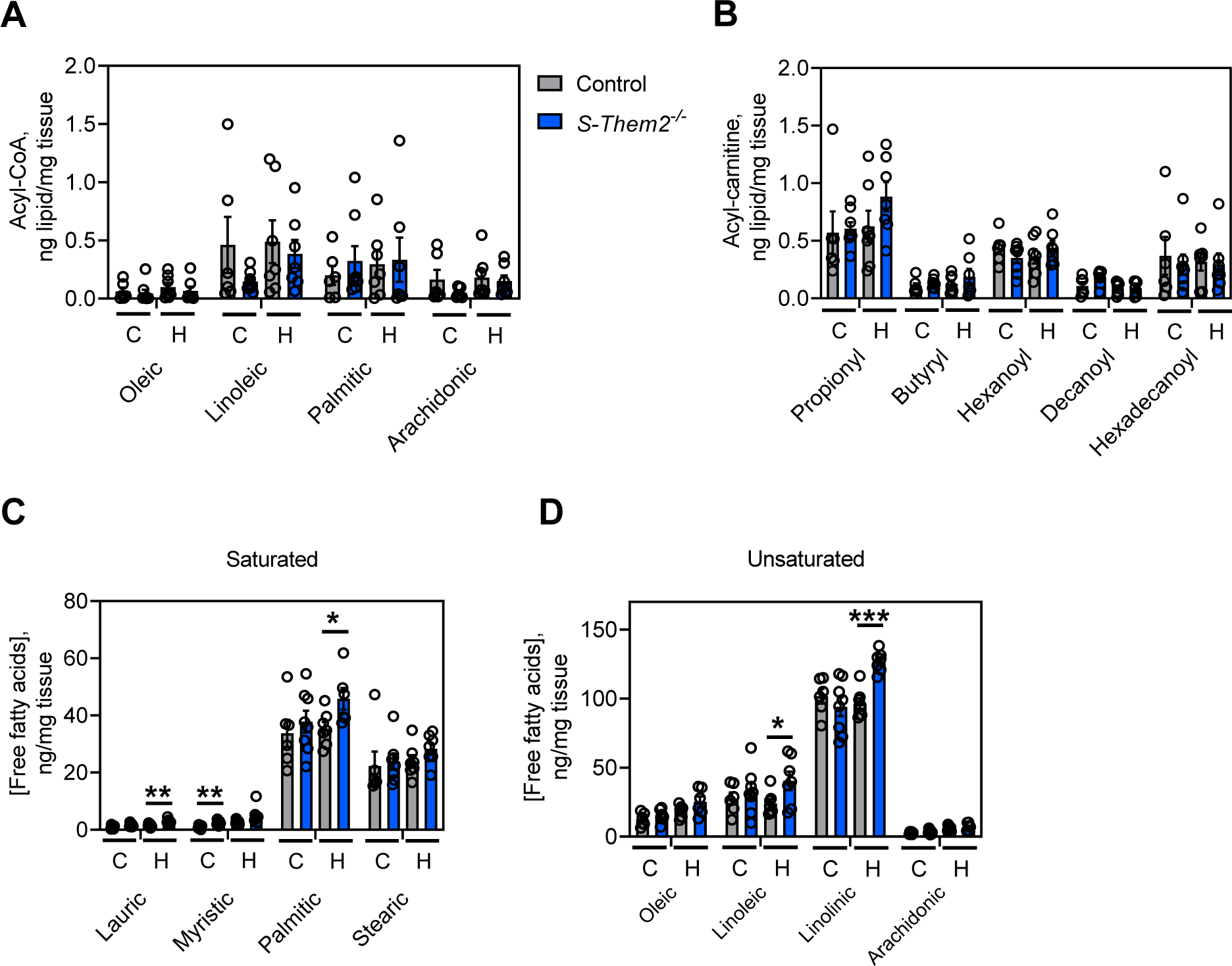
Influence of Them2 expression in skeletal muscle on molecular species of acyl-CoAs, acyl-carnitines and free fatty acids. Mice were fed chow (C) or a HFD (H) for 12 w. Molecular species of (A) fatty acyl-CoAs, (B) fatty acyl-carnitines, and (C) saturated and unsaturated long chain and short chain free fatty acids. (n = 8/ group). Statistical analyses were conducted using Student’s t test; *P < 0.05, \*\**P* < 0.01, \*\*\**P* < 0.001 vs control. Data are mean ± SEM.

**Supplemental Figure 6.**
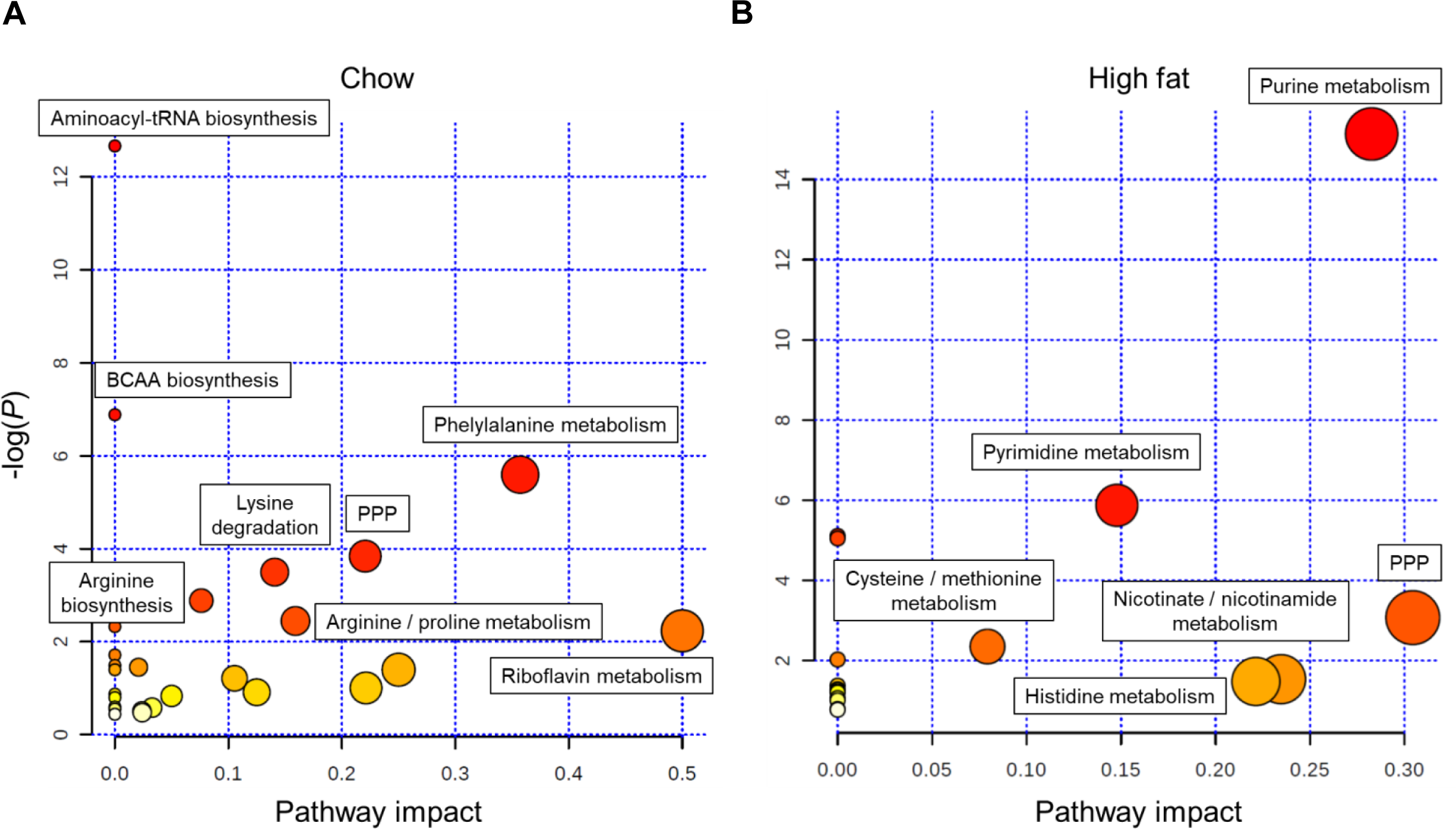
Pathway analysis of metabolomic data illustrating the influence of Them2 expression in skeletal muscle. Mice were fed (A) chow or (B) a HFD for 12 w after which gastrocnemius muscles were subjected to targeted profiling of 216 polar metabolites. Data were analyzed by MetaboAnalyst. Each node contains all the matched pathways arranged by p values (from pathway enrichment analysis) on vertical-axis, and pathway impact values (from pathway topology analysis) on horizontal-axis. The node color is based on its p value and the node radius is determined based on their pathway impact values.

**Supplemental Figure 7.**
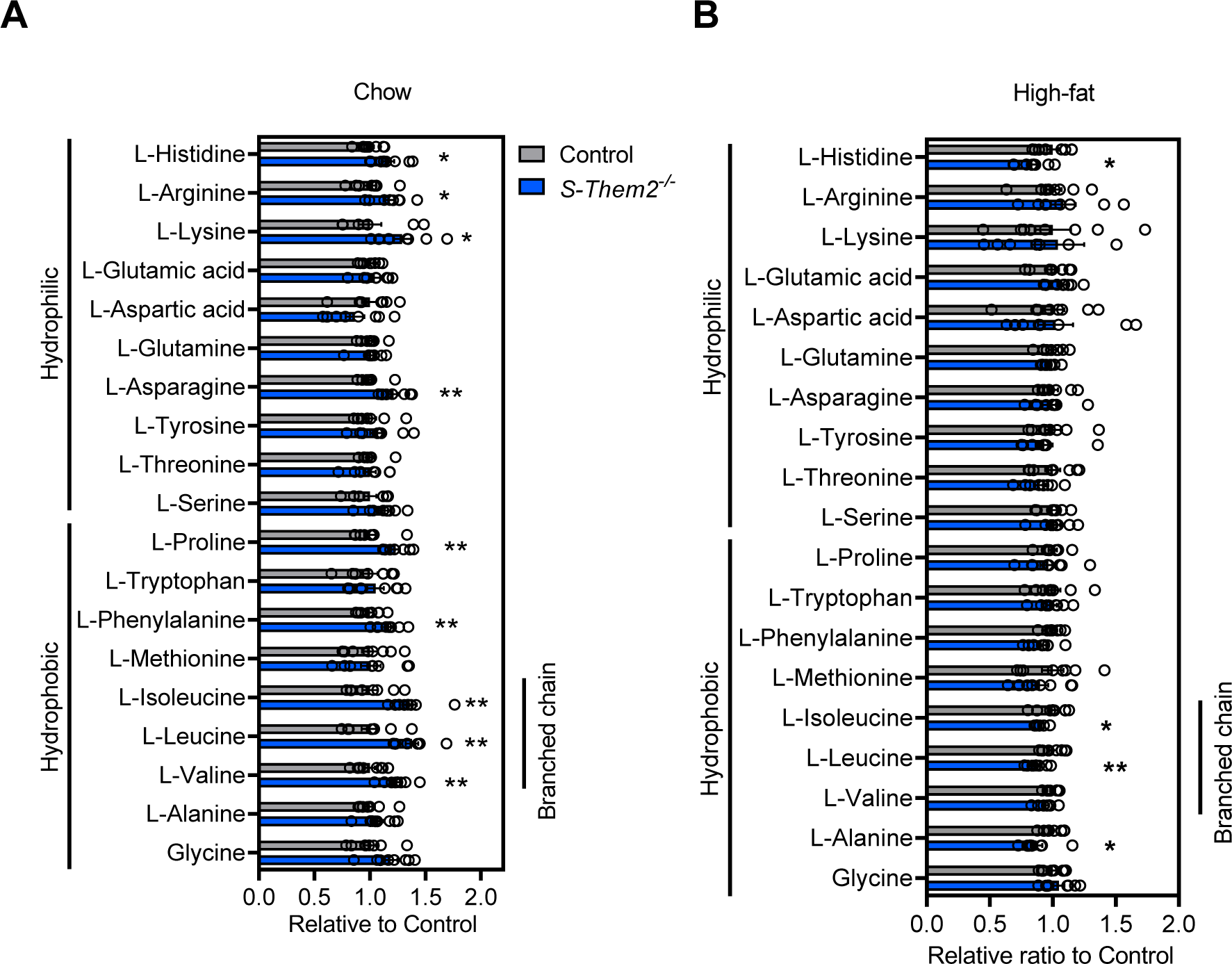
Effect of skeletal muscle Them2 expression on amino acids contents. (A,B) Relative concentrations of amino acids in gastrocnemius muscle. (n = 8/group) Statistical analyses were conducted using Student’s t test; \**P* < 0.05, \*\**P* < 0.01, \*\*\**P* < 0.001 vs Control. Data are mean ± SEM.

**Supplemental Figure 8.**
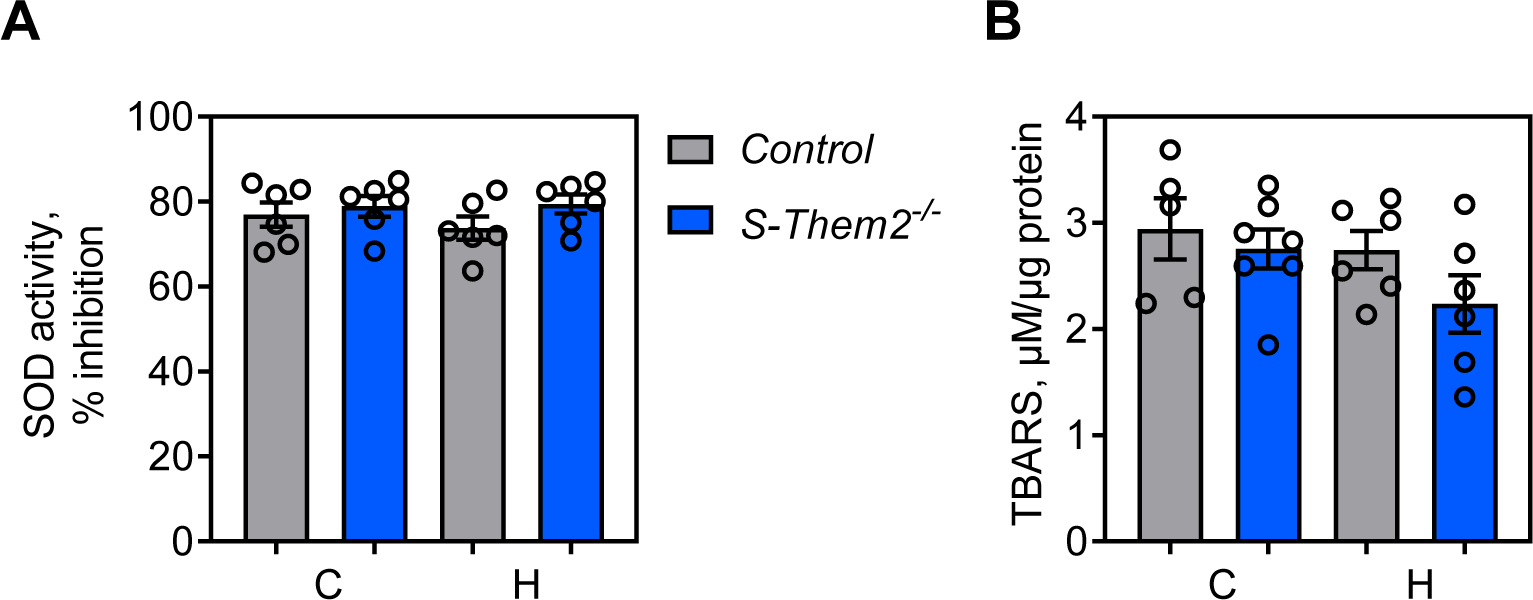
Effect of skeletal muscle Them2 expression on measures of oxidative stress. Mice were fed chow or a HFD for 12 w. (A) Superoxide dismutase (SOD) activities were assessed in skeletal muscle (n = 6/group). (B) Tissue-specific lipid peroxidation level was analyzed using a thiobarbituric acid reactive substances (TBARS) assay in skeletal muscle (n = 7/group). Statistical analyses were conducted using Student’s t test. Data are mean ± SEM

**Supplemental Figure 9.**
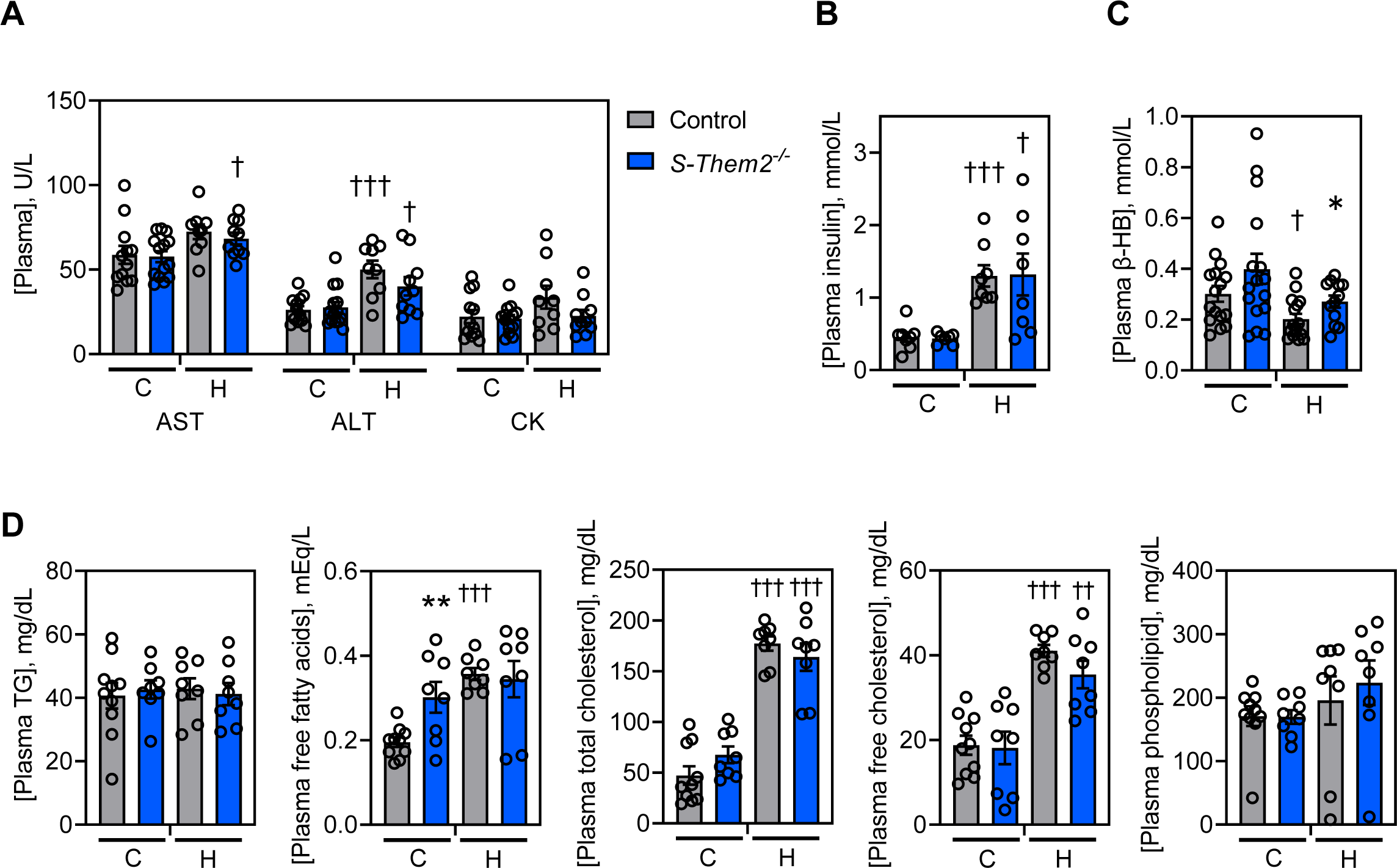
Plasma characteristics of *S-Them2^-/-^* mice. Mice were fed chow or a HFD for 12 w and the plasma was collected for (A) Activities of aspartate aminotransferase (AST), alanine aminotransferase (ALT) and creatine kinase (CK) (n = 9 – 14/group). (B) Fasting concentrations plasma insulin (n = 8/group). (C) Fasting concentrations of ß-hydroxybutyrate (b-HB/group) (n = 16). (D) Plasma concentrations of triglycerides (TG), non-esterified fatty acid (NEFA), total cholesterol (T-Cho), free cholesterol (F-Cho) and phospholipids (PL) (n = 8 – 10/group). Statistical analyses were conducted using Student’s t test; *P < 0.05, \*\**P* < 0.01 vs control. †*P*<0.05, †† *P*<0.01, ††† *P*<0.001 chow vs HFD. Data are mean ± SEM.

**Supplemental Figure 10.**
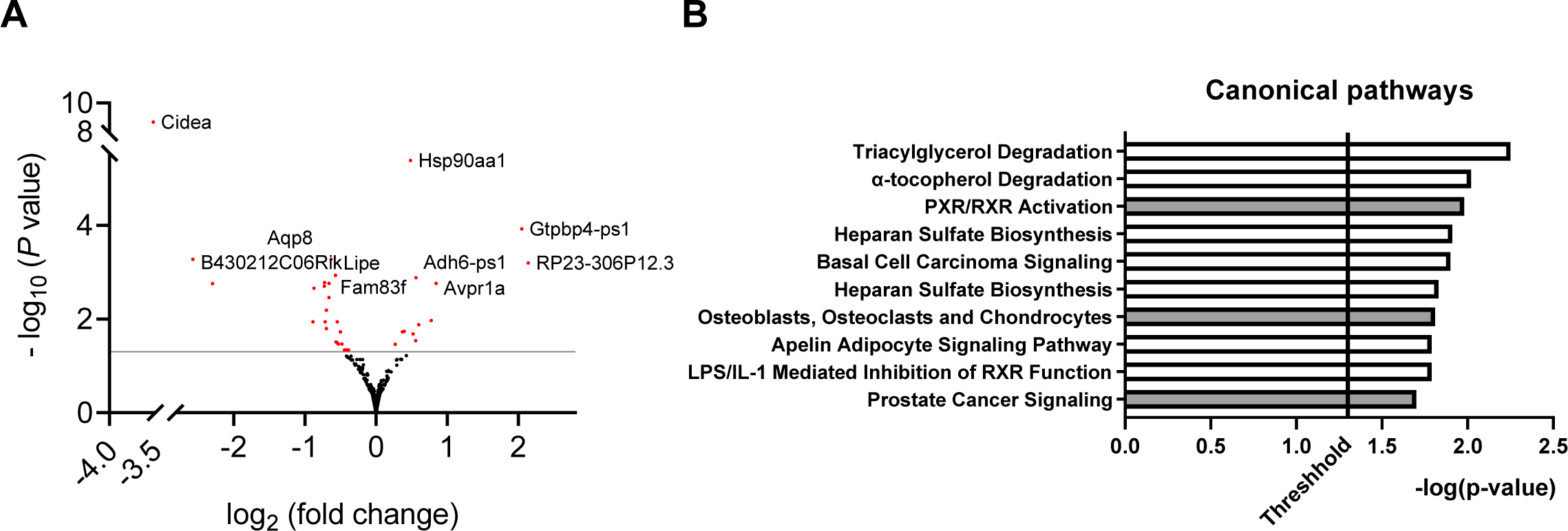
Influence of skeletal muscle Them2 expression on hepatic mRNA expression. (A) Volcano plot of RNAseq data comparing liver of control and *S-Them2*^-/-^ mice fed a HFD for 12 w. Red font represents significant changes in gene expression after adjustment for false discovery rate. The top ten genes are annotated. (B) Identification of top 10 pathways influenced by Them2 expression as assessed by Ingenuity Pathway Analysis (Qiagen) of RNAseq data. Bar color indicates predicted directionally. Z-scores, which represent directionality were calculated based on the data set’s correlation with the activated state. White indicates z-score = 0 (no activation) and gray indicates no activity pattern.

**Supplemental Table 1.**

| **Protein** | **Source** | **Reference number** |
| --- | --- | --- |
| ACC | Cell Signaling Technology (Danvers, MA, USA) | 3661 |
| Akt | Cell Signaling Technology | 9272 |
| alphaGAPDH | Novus Biologicals (Centennial, CL, USA) | NB100-56875 |
| AMPK | Cell Signaling Technology | 2532 |
| CD36 | Santa Cruz Biotechnology (Dallas, TX, USA) | 7309 |
| CPT-1 | Proteintech (Rosemont, IL, USA) | 15184-1-AP |
| HSP-90 | Santa Cruz Biotechnology | sc-25840 |
| Phospho-ACC (Ser79) | Cell Signaling Technology | 3662 |
| Phospho-Akt (Ser473) | Cell Signaling Technology | 9271 |
| Phospho-Akt (Thr308) | Cell Signaling Technology | 4056 |
| Phospho-AMPK (Thr172) | Cell Signaling Technology | 2535 |
| PCTP | Home | Shoda et al., 2001, Hepatol 33,1194-1205 |
| Them2 | Home | Kanno et al., 2007, J Biol Chem 282, 30728-30736 |

**Supplemental Table 2.**
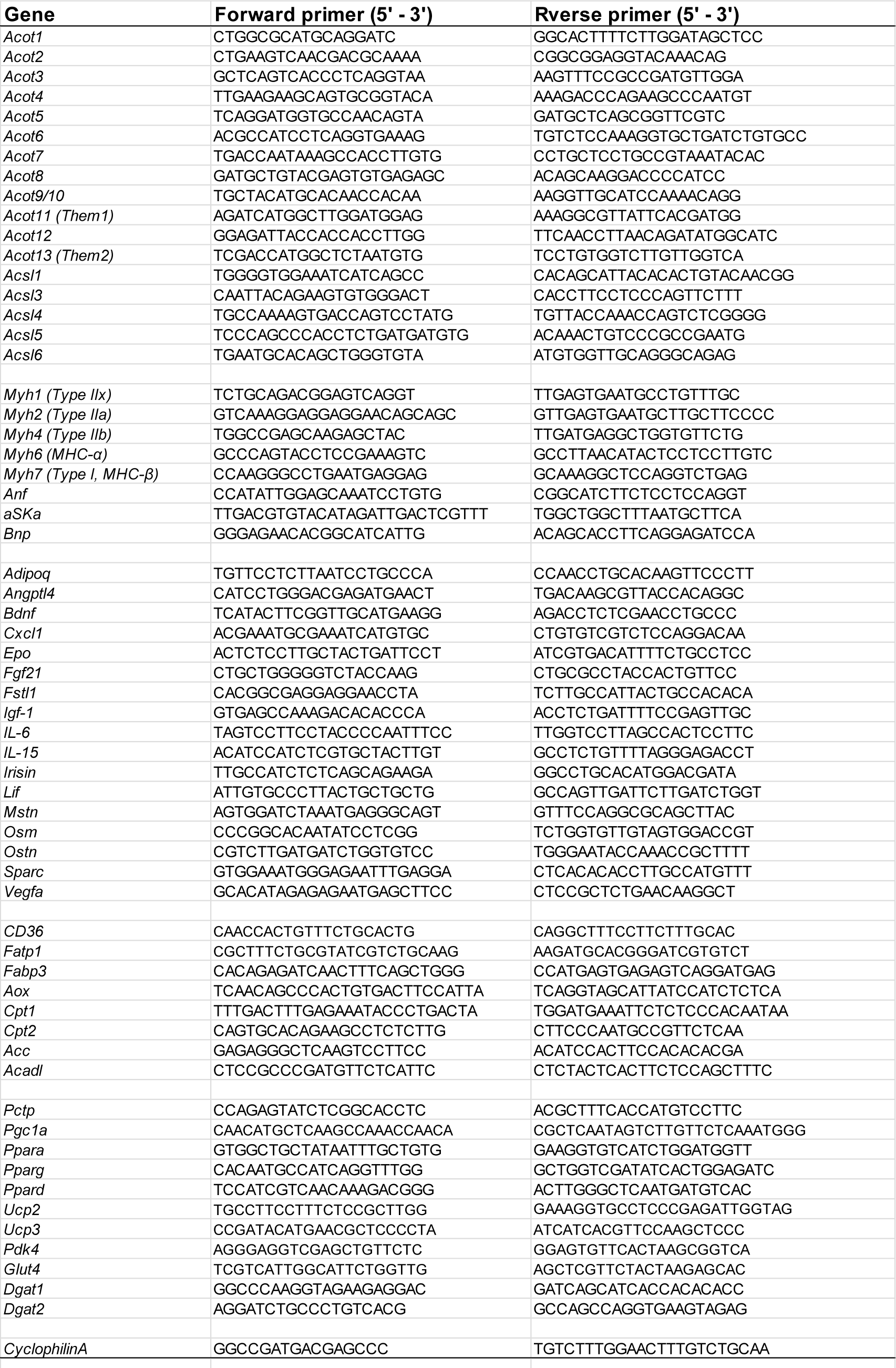

## Notes

**Financial Support:** Supported by the National Institute of Diabetes and Digestive and Kidney Diseases (R37 DK048873 and R01 DK056626 to D.E.C.); the Georgia Clinical & Translational Science Alliance of the National Institutes of Health (UL1TR002378 to E.A.O.); American Heart Association Postdoctoral Fellowship (19POST34380692 to N.I. and 18POST33990445 to M.A-B.); the American Liver Foundation Irwin M. Arias, MD Postdoctoral Research Fellowship Award American (to N.I.); the Liver Foundation NASH Fatty Liver Disease Postdoctoral Research Fellowship Award (to M.A.B.).

### Competing Interest Statement

The authors have declared no competing interest.

